# Effects of human microRNAs 100-5p, 192-5p, and 574-3p on proliferation, migration, and gene expression of human immortalized nasopharyngeal cells

**DOI:** 10.1101/2025.04.14.648665

**Authors:** Barbara G. MÜller Coan, Deilson Elgui De Oliveira

**Affiliations:** São Paulo State University (UNESP), Botucatu Medical School, Department of Pathology. Botucatu, SP, Brazil; São Paulo State University (UNESP), Institute of Biotechnology (IBTEC), Botucatu, SP, Brazil

**Keywords:** miR-100-5p, miR-192-5p, miR-574-3p, *RAB2A*, Nasopharyngeal carcinoma

## Abstract

Cancers are malignant neoplastic diseases with complex etiopathogenesis. Transformed cells acquire diverse biological properties that induce alterations in migration, invasion, and cell survival rates, favoring metastasis formation. This process is influenced by many factors, including miRNAs. MiRNAs are molecules that canonically prevent mRNA translation, influencing many cellular processes, carcinogenesis, and the progression of cancers. Nasopharyngeal carcinomas (NPCs) are aggressive cancers with high invasion rates to adjacent tissues, such as the base of the cranium, facial sinuses, and orbits, in addition to a high metastatic rate to local lymph nodes. The nonkeratinizing NPC is the most common histological form (over 95% of cases) and is highly related to Epstein-Barr virus (EBV) infection, notably when undifferentiated. In a previous study, we observed that immortalized nasopharyngeal cells NP69^SV40T^ transfected to express the EBV Latent Membrane Protein 1 (LMP1) from the M81 strain hyperregulate human microRNAs miR-100-5p, miR-192-5p, and miR-574-3p, compared to NP69^SV40T^ cells transfected with LMP1 from the EBV strain B95.8A. In this study, we aimed to assess the putative effects of each of the identified microRNAs on the viability, migration, and proliferation of NP69^SV40T^ cells cultivated in vitro. Therefore, target prediction and pathway enrichment analysis were performed to infer which pathways could be modulated by each miRNA, followed by expression analysis of selected predicted target genes. Additionally, NP69^SV40T^ cells were transfected with miR-100-5p, miR-192-5p, or miR-574-3p mimics to analyze cell behavior. After their transfection with the miR-192-5p mimic, we observed a 40% decrease in *RAB2A* transcript levels and a 43% reduction in cell migration. During the pathway enrichment analysis, miR-192-5p showed predicted involvement in pathways such as MAPK, GPCRs, TKR, and TGF-β. NP69^SV40T^ cells transfected with miR-100-5p or miR-574-3p reduced transcriptional levels of their targets FZD8 and STC1, respectively. However, in the performed cellular assays, they did not alter cell behavior. The results demonstrate that miR-192-5p acts as a tumor suppressor, elucidating part of its function in nasopharyngeal cells. This highlights its importance in the NPC context and the necessity of further studies on this miRNA.

## Introduction

Cancers are a group of neoplastic diseases with complex etiology. In 2024, the Global Cancer Observatory (GCO) accounted for almost 20 million new cancer cases and 9.7 million deaths directly attributed to these diseases ^**[#1 - Ervik M et al., 2024 | #2 - Sung et al., 2021 | #3 - Thun et al., 2010]**^. Nasopharyngeal carcinoma (NPC) is a highly aggressive cancer strongly associated with infection by the Epstein-Barr virus (EBV)^**[#4 - Liu et al., 2022]**^, as well as environmental (e.g., exposure to nitrosamides, diet) and genetic factors (e.g., HLA genes on chromosome 6p21) ^**[ #5 - Chen et al., 2019]**^. In 2024, 120,434 new cases and over 73,000 associated deaths were related estimated worldwide for NPC, which is endemic in the Southeast Asia, notably in China ^**[#2 - Sung et al., 2021]**^. NPC invades tissues adjacent to the primary tumor, such as the base of the cranium, facial sinuses, and orbits. Furthermore, at diagnosis the disease manifests with metastasis in regional lymph nodes in about 50% of cases. Disseminated disease usually occurs within 3 years after initial presentation, and the most common metastasis sites are bones, lungs, and liver ^**[#5 - Chenet al., 2019 | #6 - Su et al., 2022 | #7 - Thompson, 2007]**^.

Overall, metastatic dissemination requires that malignant cells acquire particular properties that enable them to colonize tissues distinct from the primary site and is related to poor cancer prognosis, increased morbidity, and mortality rates ^**[#8 - Kiri and Ryba, 2024 | #9 - Tang et al., 2020]**^. The metastasis process is influenced by non-coding microRNAs (miRNAs) ^**[#10 - Choi and Lee, 2013 | #6 - Su et al., 2022]**^, small non-coding transcripts that negatively regulate mRNA translation (most commonly, canonical activity). Alteration in miRNA expression in cancers has been widely described, and these molecules can act as oncomirs, promoting cancer development (targeting tumor suppressor genes), as well as tumor suppressor miRNAs (targeting oncogenes), or even with dual activity, in a context-dependent manner, for instance ^**[#11 - Ali Syeda et al., 2020 | #12 - Looi et al., 2023]**^.

In a previous study ^**[#13 - Müller Coan et al., 2022]**^, we identified the human miRNAs 100-5p, 192-5p, and 574-3p as deregulated miRNAs in immortalized nasopharyngeal cells transfected with the Epstein-Barr virus (EBV) oncoprotein LMP1 from two viral variants (M81 and B95.8A). We found that miR-192-5p was 2.7-fold downregulated in the NP69^SV40T^ (RRID:CVCL_F755) immortalized nasopharyngeal cells transfected with the EBV LMP1 oncoprotein derived from the prototype viral strain B95.8A (originated from a Burkitt lymphoma, used as a control) compared to cells not expressing LMP1, and miRNAs 100-5p, 192-5p, and 574-3p were upregulated (1.8, 1.7, and 2-fold, respectively) comparing EBV LMP1 derived from the viral strain M81 (originated from NPC) compared to the B95.8A strain. However, no differences in these three miRNAs were found when comparing EBV LMP1 from M81 to cells not expressing LMP1.

The obtained result suggests a putative pathogenetic role of the indicated miRNAs in the pathogenesis of NPC, prompting further investigation of their biological roles on human cells of the nasopharyngeal epithelium. Therefore, in this study we aimed to investigate how the microRNAs miR-100-5p, miR-192-5p, and miR-574-3p impact the behavior of human immortalized nasopharyngeal cells (NP69) cultivated *in vitro*, transfected with the respective miRNA mimics, regarding gene expression, migration, and proliferation. as will be detailed further here, we found that miR-192-5p could downregulate the transcript levels of its target gene, *RAB2A*, by 40% and decreased migration of NP69 cells in 43%. Despite miR-100-5p and miR-574-3p causing a decrease in its target genes, *FZD8* and *STC1* respectively, they could not induce changes in cell behavior. This indicates that miR-192-5p is a good candidate for further molecular characterization and cell behavior effects *in vitro* and *in vivo* studies in nasopharyngeal carcinoma.

## Material and Methods

### Cell culture

This study was conducted with the immortalized human nasopharyngeal cell NP69^SV40T^ ([RRID:CVCL_F755](https://www.cellosaurus.org/CVCL_F755)), herein indicated as NP69. This cell line was generated by the research group headed by Professor George Sai-Wah Tsao (Hong Kong University, China), and it was obtained from Dr. Ethel Cesarman (Cornell University, NY, USA). The cells were cultivated with the Keratinocyte SFM medium (Thermo Fisher Scientific, Waltham, M, USA) supplemented with 5% Fetal Bovine Serum (FBS), EGF and BPE (according to the manufacturer’s instructions), and 0.4% gentamicin for microbiological control. The genetic identity of the cell line was confirmed in our lab by short tandem repeats (STRs) analysis using the GenePrint 10 (Promega, Madison, WI, USA), according to the manufacturer’s instructions. The obtained cell profile matched the profile deposited for NP69 cells (TH01: 7; D21S11: 31; D5S818: 11; D13S317: 10, 12; D7S820: 11; D16S539: 11, 12; CSF1PO: 12, 13; AMEL: X, Y; vWA: 16, 19; TPOX: 11). Cell cultures were also confirmed to be free of *mycoplasma* contamination using a PCR-based protocol ^**[#14 - Uemori et al., 1992]**^.

### miRNA Target Prediction and Pathway Enrichment Analysis

For each miRNA investigated, we performed *in silico* target prediction (Supplementary Material, Figure S1) and pathway enrichment analysis to identify predicted transcripts that each miRNA could be targeting and, consequently, the pathways and cellular changes could be expected upon miRNA ectopic expression using miRNA mimics. The target prediction analysis was performed using the online tool mirDIP ^**[#15 - Shirdel et al., 2011 | #16 - Tokar et al., 2018]**^ (Supplementary Material, Table S1) and Pathway Enrichment Analysis was performed using the ReactomeFIViz ^**[#17 - Croft et al., 2014]**^ plugin in Cytoscape ^**[#18 - Shannon et al., 2003]**^ (Supplementary Material, Table S2). The analysis of complementarity between the miRNA seed region and the 3’ UTR region of target genes was performed using the 3’ UTR sequence Database ^**[#19-Grillo et al., 2010]**^, and the *in silico* pairing was confirmed using the NCBI Blast online tool ^**[#20 - Altschul et al.,1990 | #21-Boratyn et al., 2013]**^.

### Cellular transfection of miRNA mimetics and assessment of miRNA expression

Before each functional experiment using miRNA mimics, NP69 cells were maintained under serum starvation (medium without FBS) for 24h, counted and plated into a 12- or 24-well plate. After 24 h of incubation, the cells were transfected using Lipofectamine™ RNAiMAX Transfection Reagent (Thermo Fisher) with 10nM of MISSION® miRNA Mimic (Sigma-Aldrich, St Louis, MO, USA), following manufacturer’s instructions. For all experiments, the cells were incubated with the transfection reagent for 24 h, dissociated with trypsin, counted using Trypan blue (used for viability assessment), and subjected to the specific downstream experiment (mRNA analysis, migration, or proliferation assays).

To analyze the relative amount of selected miRNA target transcripts, total RNA was extracted from NP69 cells transiently transfected (miR-100-5p, miR-192-5p, or miR-574-3p mimic) using the TRIzol™ Reagent (Thermo Fisher), according to the manufacturer’s instructions. After extraction and quantification, the RNA integrity was indirectly assessed by inspection of ribosomal bands and absence of smears in 1% agarose gels after total RNA electrophoresis. Following, the cDNA synthesis was performed to analyze miRNA levels (Supplementary Material, Figure S2) and selected miRNA targets (Figure 3). MiRNA reverse transcription was performed using High-Capacity cDNA Reverse Transcription Kit (Thermo Fisher) with random primers supplied with the miRNA qPCR kits (Canopy Biosciences). The High-Capacity cDNA Reverse Transcription Kit (Thermo Fisher) was also used to produce cDNA subjected to analysis of the expression of predicted miRNA target transcripts. All protocols were performed following the manufacturer’s recommendations. The reverse transcription real-time quantitative PCR (RT-qPCR) assays to assess miRNA and gene expression were performed using the GoTaq® qPCR system in AriaMx Real-Time PCR (qPCR) Instrument (Agilent) and 7500 Real-Time PCR (Applied Biosystems). All RT-qPCR experiments were performed in technical duplicates and biological triplicates, using 2 reference genes. Primers and reaction components are in Supplementary Material, Table S3.

### Estimation of cell proliferation and migration rates in vitro

To estimate the cell proliferation rates, NP69 cells transfected with control (cells incubated with the transfection reagent only), miR-100-5p, miR-192-5p, or miR-574-3p were dissociated 24 h post-transfection, counted, and distributed in duplicate in five 96 well plates. These cells were incubated in complete media and assessed in 24 h intervals at days 0, 1, 2, 3, and 4 post-transfection using the colorimetric assay CellTiter 96® Aqueous One Solution (Promega), following the manufacturer’s protocol, with absorbance readings at 490 nm performed with the spectrometer reader device (Bio-Rad Model 680).

For cell migration, NP69 cells were dissociated 24 h post-transfection, counted, and inoculated (1 x 10^5^ cells) in the inner chamber of Transwell inserts (8µm pores size) filled with serum-free media and adjusted into wells filled with complete media in a 24-well plate. After an 18h incubation, the inner side of the membrane was scraped with a cotton swab (to remove cells that did not migrate), fixed with 70% ethanol for 10 min, and stained with crystal violet for 15 min. ImageJ software ^**[#22 - Schneider et al., 2012]**^ was used to count 25% of the total area of the outer membrane of the insert, considering migrating cells. The assay was performed with technical duplicates and three independent experiments were performed (biological triplicates).

## Results

### In silico analysis of miR-100-5p, miR-192-5p and miR-574-3p showed unique deregulated pathways and target genes

We performed an *in silico* target prediction analysis to identify putative targets for miR-100-5p, miR-192-5p, and miR-574-3p, along with pathway enrichment analysis to predict potential changes in cell behavior in which the investigated miRNAs could be implicated when expressed in the human immortalized nasopharyngeal cell line NP69.

Accounting only for the top 1% predicted targets from the three miRNAs, there were 82, 545, and 10 selected genes for miR-100-5p, miR-192-5p, and miR-574-3p, respectively (Supplementary Material, Table S1 and Figure S1). The pathway enrichment analysis (Supplementary Material, Table S2) provided several insights on behavioral changes induced by each of the miRNAs. The results are shown in Figure 1.

**Figure 1.**
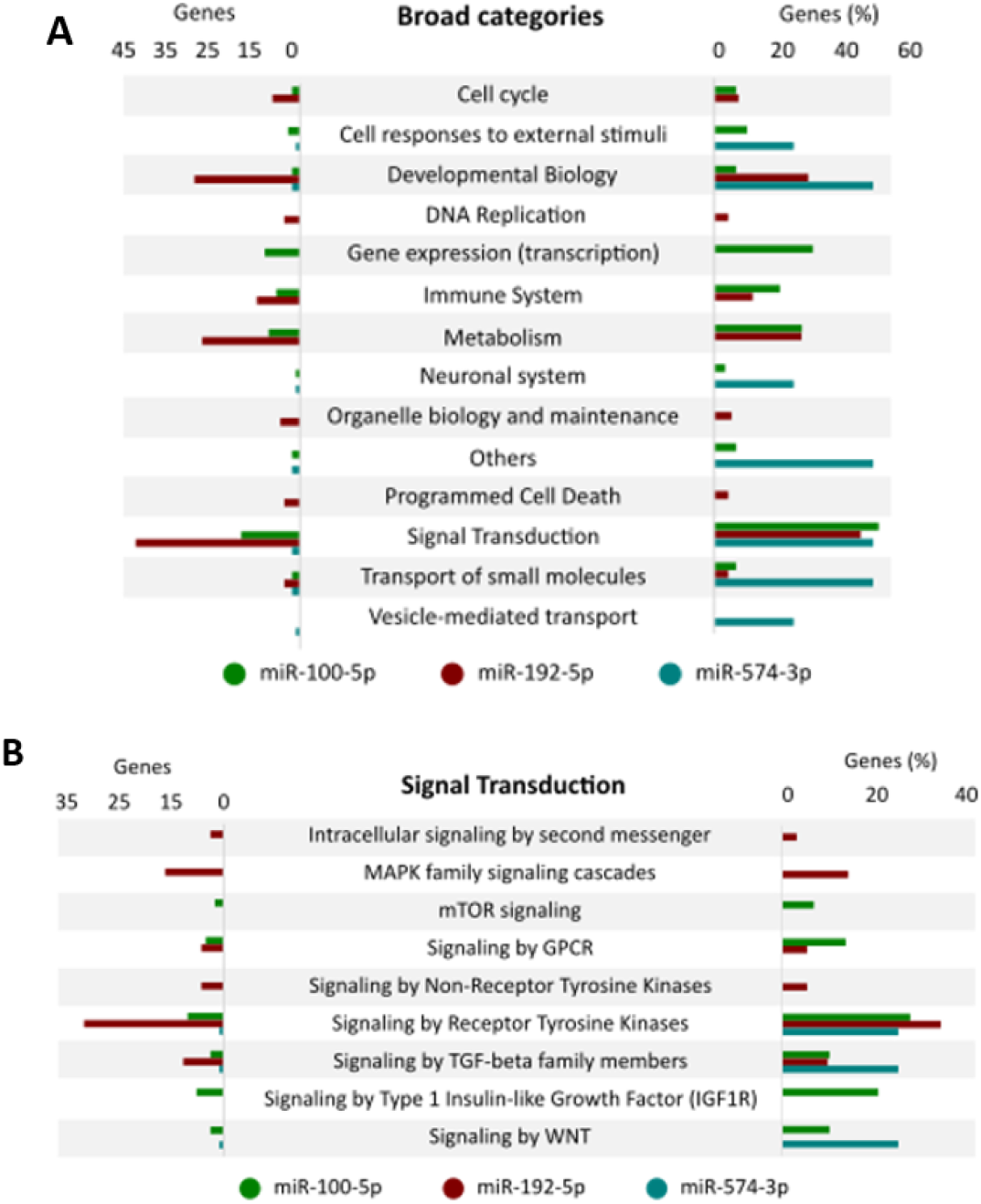
Pathway enrichment analysis for high-score target genes identified for miR-100-5p, miR-192-5p, and miR-574-3p, selected from a previous work ^[#23 - Müller-Coan and Elgui de Oliveira, 2016 p. 1]^ . (A) Broad cellular pathways categories and their respective gene hits. (B) Gene hits obtained for pathways within the Signal Transduction category.

Briefly, the miR-100-5p was predicted to be involved in modulation of gene expression, immune system, metabolism, and signal transduction pathways (Figure 1A), which comprises more than 50% of target genes predicted. Some known pathways that are critical in cancer biology were predicted to be regulated by this miRNA, such as mTOR, GPCR, TGF-β, IGF1R, FGFR3b, and WNT (Figure 1B and Supplementary Material, Table S2). Regarding miR-192-5p, the analysis predicted its activity regulating biological development pathways, metabolic routes, and signal transduction pathways (Figure 1A). Over 40% of the predicted genes were involved in MAPK, GPCR, TKR, and TGF-β pathways, and approximately 35% of them are related to tyrosine-kinase receptors (TKRs) pathways, such as EGFR, ERBB2, FGFR, PDGF, IGF1R e NGF (Figure 1B and Supplementary Material, Table S2). Finally, miR-574-3p had predicted effects on cellular responses to external stimuli, biological development, vesicle-mediated transport and biogenesis, cellular response to hypoxia, as well as signal transduction pathways (Figure 1A and Supplementary Material, Table S2). MiR-574-3p has a smaller number of predicted target genes selected for analysis, but the small number does not prevent further studies. During the pathway enrichment analysis, miR-574-3p predicted targets were related to important pathways in cancer, such as TGF-β, WNT, HIF, and TKR signaling (Figure 1B and Supplementary Material, Table S2).

Overall, all three of the miRNAs investigated were predicted to be involved in developmental biology, transport of small molecules, and in some specific cellular pathways, such as Bone Morphogenetic Protein signaling (BMP), TGF-β, and TKR (Figure 1B and Supplementary Material, Table S2). In common, miR-100-5p and miR-192-5p were predicted to regulate the immune system, cell cycle, and the GPCR pathway, while miR-192-5p and miR-574-3p were predicted to modulate signaling events driven by the adhesion molecule L1CAM and NODAL (Supplementary Material, Table S2). Finally, miR-100-5p and miR-574-3p showed up as regulators of VLDLR internalization and degradation, VLDL interactions, and Lipoprotein metabolism (Supplementary Material, Table S2).

These results obtained after *in silico* analysis of predicted targets of the miRNAs investigated also suggest several cell behavior modifications individually induced by each of the miRNAs, such as mTOR signaling by miR-100-5p, GPCR signaling by miR-192-5p, and vesicle biogenesis and transport by miR-574-3p. Considering all three miRNAs, they were predicted to regulate pathways involved in proliferation, migration, and apoptosis. In conclusion, all three miRNAs could alter cell behavior through different mechanisms, including in the NP69 transfected with its mimics.

### Effects of treatment of NP69 cells with mimetics for miR-100-5p, miR-192-5p, and miR-574-3p in selected gene targets

To validate the results of the *in silico* analysis performed, the expression of transcripts for selected target genes predicted for each of the investigated miRNAs was assessed by qPCR. As shown in Figure 2 (Panels A to C), all the selected gene targets for miRs 100-5p, 192-5p, and 574-3p had complementarity in at least one site at the target 3’-UTR region.

**Figure 2.**
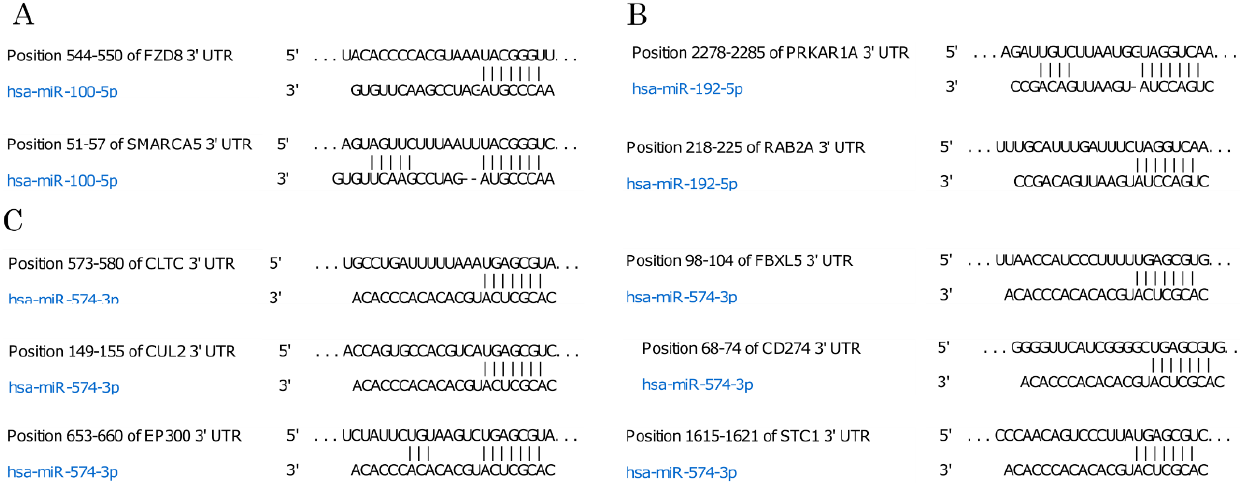
Complementarity analysis of whole miRNA sequence and the 3’UTR region of the respective targets for miR-100-5p, miR-192-5p, and miR-574-3p. Panel A shows the complementarity between miR-100-5p and the 3’-UTR sequence from FZD8 and SMARCA5 transcripts. Panel B shows the complementarity between miR-192-5p and the 3’-UTR region from PRKAR1A and RAB2A transcript. Panel C shows the complementarity between miR-574-3p and the 3’-UTR sequence from CUL2, CLTC, EP300, FBXL5, PD-L1, and STC1 transcripts. Matching nucleotides are highlighted in bold, and the miRNA seed region is highlighted in green. Image obtained from TargetScan 7 ^[#24 - Agarwal et al., n.d.]^.

For miR-100-5p, the mRNA levels were assessed for the genes *FZD8* (NCBI Gene ID: 8325), which encodes a frizzled receptor for the WNT pathway, and *SMARCA5* (NCBI Gene ID: 8467), encoding a helicase with nucleosome remodeling activity. NP69 cells transfected with the miR-100-5p mimic showed a 40% decrease in *FZD8* levels, and no significant change for *SMARCA5* (Figure 3A).

**Figure 3.**
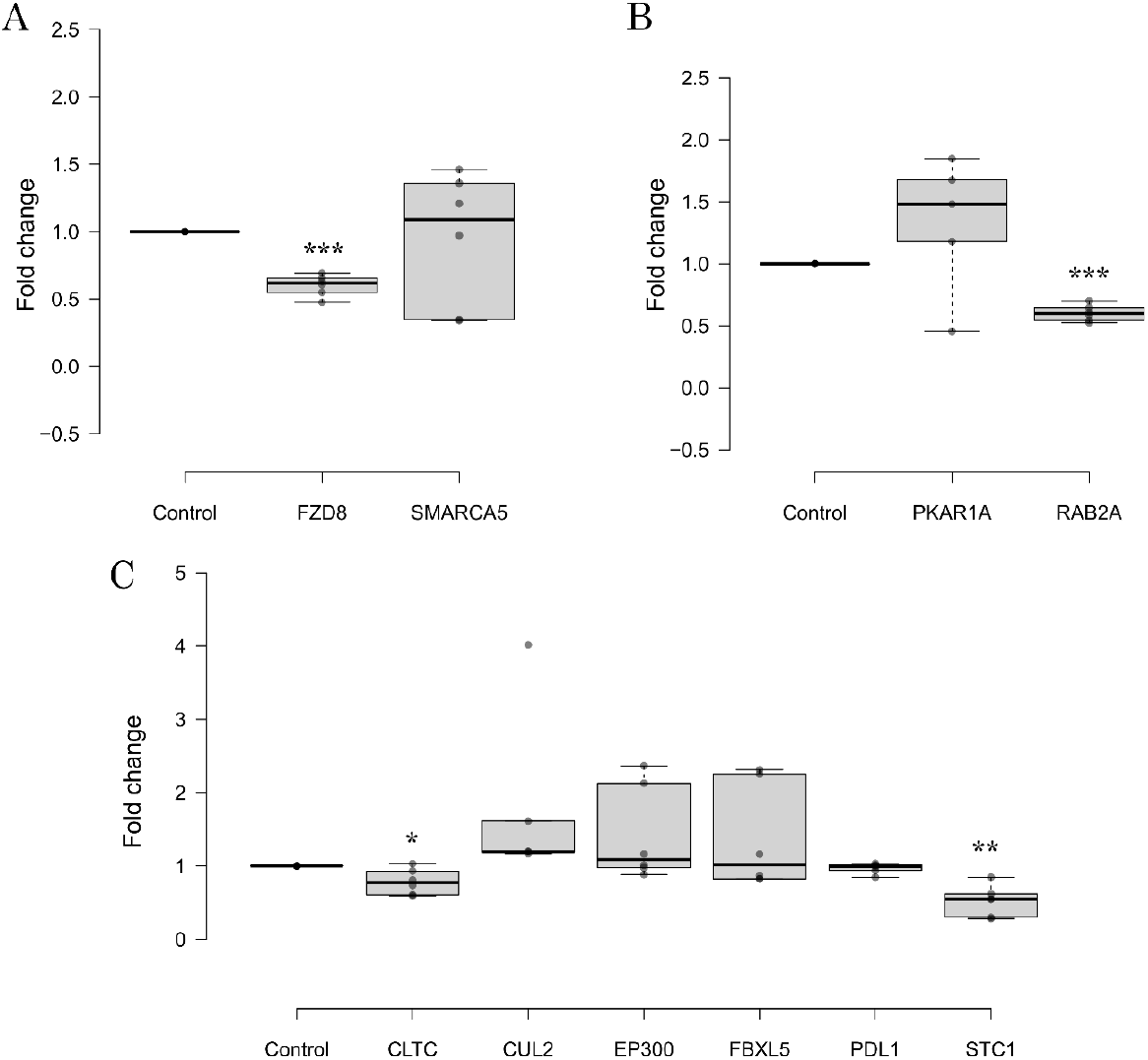
Assessment of mRNA expression of gene targets predicted for (A) miR-100-5p (*FZD8* and *SMARCA5*), (B) miR-192-5p (*PRKAR1A* and *RAB2A*), and (C) miR-574-3p (*CUL2, CLTC, EP300, FBXL5, PDL1* and *STC1*) in NP69 cells transfected with 10nM of the respective microRNA mimic. A 40% reduction in mRNA expression levels was found for *FZD8* and *RAB2A* in cells treated with mimics for miR-100-5p and miR-192-5p, respectively. Furthermore, cells treated with the miR-574-3p mimic showed a reduction of 48% on the transcriptional levels of *STC1*. No significant changes were observed for other conditions/targets. * p<0.05; ** p<0,005 and ***p<0.002.

For miR-192-5p, the mRNA levels were assessed for the genes *PKRAR1A* (NCBI Gene ID: 5573) which encodes for a protein kinase CAMP-Dependent, and *RAB2A* (NCBI Gene ID: 5862), a small GTPase member of the RAS family. NP69 cells transfected with the miR-192-5p mimic showed a 40% decrease in *RAB2A* levels, and no significant change for *PRKAR1A* (Figure 3B).

For miR-574-3p, mRNA levels were assessed for the genes *CLTC* (NCBI Gene ID: 1213), which encodes for a clathrin, a component of the cytoplasmic face from organelles involved in the formation of coated vesicles; *CUL2* (NCBI Gene ID: 8453), which encodes a cullin protein, a core component of multiple cullin-RING-based ECS (ElonginB/C-CUL2/5-SOCS-box protein) related to hypoxia; *FBXL5* (NCBI Gene ID: 26234), a member of the F-box protein family involved in protein ubiquitination; and *STC1* (NCBI Gene ID: 6781), a glycoprotein with autocrine and paracrine functions involved in cell metabolism and calcium/phosphate homeostasis. Other targets genes predicted to be not included in the top 1% list were also included due to their relevance in the context of cancer. The transcriptional levels of these genes were also assessed: *EP300* (NCBI Gene ID: 2033), a histone acetyltransferase that regulates transcription via chromatin remodeling, and *CD274* (also called *PDL1* – NCBI Gene ID: 29126), an immune inhibitory receptor ligand. NP69 cells transfected with the miR-574-3p mimic showed a 48% decrease in *STC1* levels, and no significant change for *CLTC, CUL2, EP300, FBXL5* and *PDL1* (*CD274*). (Figure 3C).

### MiR-192-5p mimic treatment decreases cell migration but does not induce alterations in cell proliferation or viability in vitro

To investigate *in vitro* effects induced by the treatment with miR-100-5p, miR-192-5p, or miR-574-3p (Supplementary Material, Figure S2), 24h post-transfection with the miRNA mimetics, the NP69 cells were subjected to assays to assess cell viability (trypan blue dye exclusion assay; Figure 4), cell proliferation rates (colorimetric assay with cell titer; Figure 5), and cell migration (Transwell assay; Figure 6).

**Figure 4.**
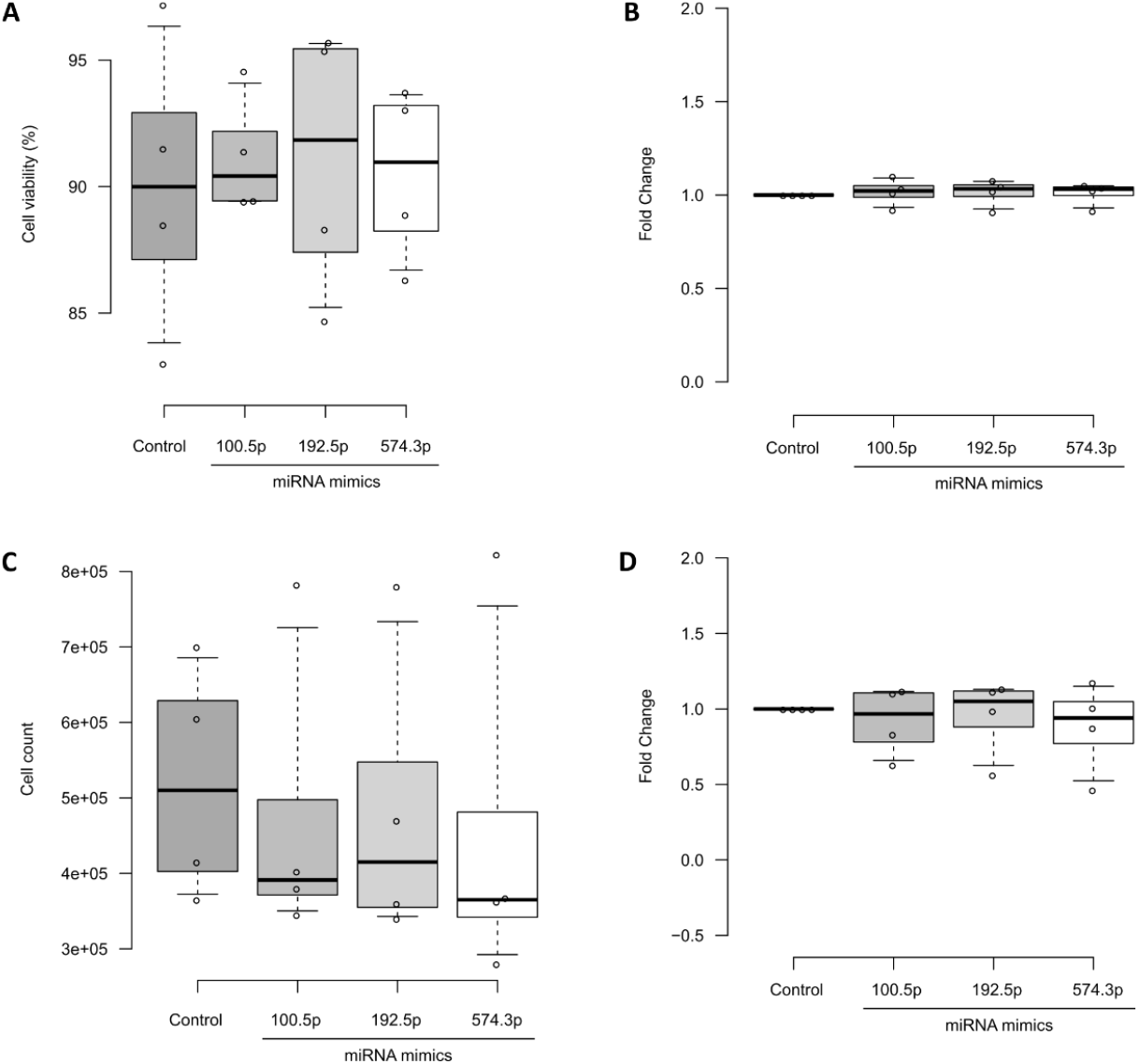
Assessment of (A) total cell count, (B) cell count relative to control, (C) cell viability and (D) cell viability relative to control in NP69 cells transfected with 10nM of the miR-100-5p, miR-192-5p or miR-574-3p mimic. Cell count and viability were performed using trypan blue stain and Neubauer chamber using trypan blue stain. (A) shows the total percentage of cell viability; (B) shows the cell viability relative to control; © shows the total number of counted cells; and (D) shows the number of counted cells relative to the control. No significant changes were observed for any of the conditions.

**Figure 5.**
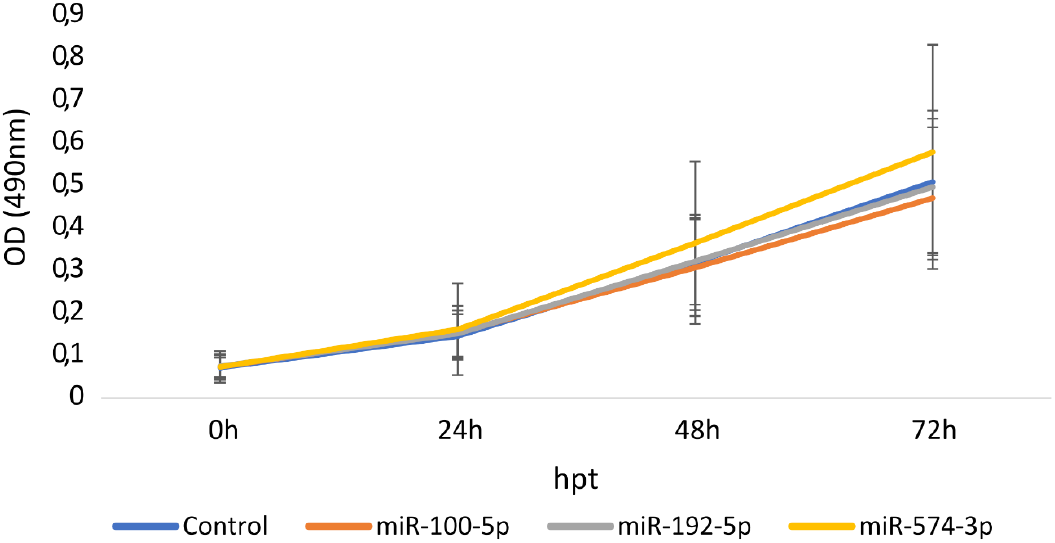
Proliferation rate of NP69 cells transfected with mimetics for miRs 100-5p (orange), 192-5p (grey), miR-574-3p (yellow), or the control (blue) using an MTT-based assay (Promega CellTiter 96® AQueous One Solution reagent). Readings were started 24h post-transfection (0h). No significant changes were observed for any of the conditions.

**Figure 6.**
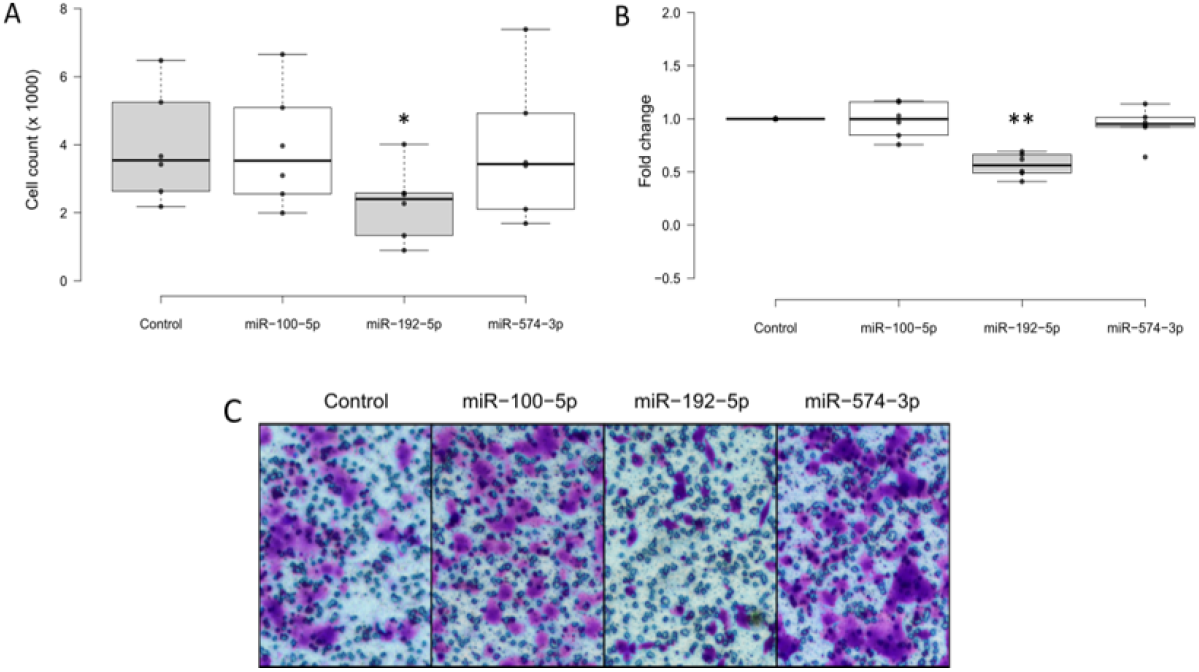
Migration analysis using transwell assay of NP69 cells transfected with the mimic for miR-100-5p, miR-192-5p, miR-574-3p or the control (transfection reagent only). (A) shows the total number of cells that migrated through the transwell membrane; (B) shows shows the relative number of migrating cells compared to the control; (C) shows a representative image of migrating cells. A 43 % reduction in migration rates was found in cells treated with miR-192-5p mimic. No significant changes were observed for miR-100-5p or miR-574-3p mimic treatment. * p < 0.001; ** p < 0.0001.

Treatment with miR-192-5p caused a 43% reduction in the number of migrating NP69 cells compared to the control (Figure 6) and no statistically significant differences were found in viability (Figure 4A and B), cell count (Figure 4C and D), or proliferation (Figure 5). Regarding treatments with miR-100-5p or miR-574-3p mimic in NP69 cells, no statistically significant differences were found in cell viability (Figure 4A and B), cell count (Figure 4C and D), proliferation (Figure 5), or migration (Figure 6).

## Discussion

MiRNAs have been described as important molecules involved in cancer development and progression. MiR-100-5p, miR-192-5p, and miR-574-3p have been previously related to involvement in several cellular pathways and to modulating malignant cell behavior, such as migration, invasion, proliferation, EMT, and metastasis rates.

MiR-100-5p has a tumor suppressor function, and our work shows that it can potentially modulate several cellular pathways important in cancer, such as mTOR, IGF1R, and TKR. Consistent with our prediction analysis, miR-100-5p has been already described to regulate mTOR in cardiac hypertrophy ^**[#25 - Zeng et al., 2021]**^. Additionally, it is downregulated in chordoma and nasopharyngeal carcinoma samples, and, when upregulated, reduced migration, invasion, and proliferation of those cells through IGF1R pathway ^**[#26 - Sun et al., 2018 | #27 - Zhang et al., 2020]**^. Despite its primary role as a tumor suppressor, silencing of miR-100-5p by FOXA1 inhibited the malignant behavior of nasopharyngeal carcinoma cells. In contrast, upregulation of miR-100-5p induced malignant features in NPC cells, such as increased migration and colony formation, through the inhibition of RASGRP3 or FOXN3 translation, thus acting as an oncomiR ^**[#28 - Peng et al., 2020]**^.

Transfection of NP69 cells with miR-100-5p mimic showed a 40% reduction in *FZD8* transcripts, and the correlation with miR-100-5p as a target is yet unpublished. The FZD8 protein is known to be involved in several key signaling cascades affected in cancers ^**[#29 - Le et al., 2015]**^, including the TGF-β and Wnt pathways, predicted by the *in silico* analysis performed in this study (Figure 1B). TGF-β is known to promote *FZD8* expression, leading to non-canonical activation of Wnt via its WNT-5b ligand ^**[#30 - Spanjer et al., 2016]**^. Along with *FZD5, FZD8* is essential for Wnt activation ^**[#31 - Alok et al., 2017]**^ and is being explored as a therapeutic target to inhibit this pathway ^**[#29 - Le et al., 2015]**^. The activation of the Wnt pathway (either canonical or non-canonical) by *FZD8* was reported to increase cell proliferation rates and metastasis in renal cell carcinomas ^**[#32 - Yang et al., 2017]**^, as well as the frequency of bone metastasis and acquisition of cancer stem cell (CSC) properties in prostate cancers ^**[#33 - Li et al., 2017]**^.

The crosstalk between TGF-β, miR-100, and Wnt signaling has been described in osteocytes. In this context, TGF-β suppresses miR-100, which antagonizes the Wnt pathway, mimicking fluid shear stress-induced Wnt activation and regulating the mechanosensitive function of osteocytes ^**[#34 - Dole et al., 2021]**^. Similarly, in thyroid cancer cells, miR-100-5p inhibits the Wnt/β-catenin pathway by targeting FZD8, leading to reduced cell proliferation and induced apoptosis ^**[#35 - Ma and Han, 2022]**^.

The results reported here provide relevant insights into the effects of upregulation of miR-192-5p in nasopharyngeal cells. The pathway prediction analysis showed that over 40% of the predicted genes were involved in MAPK, GPCRs, TKR, and TGF-β pathways, and approximately 35% of them are related to tyrosine-kinase receptors (TKRs) pathways, such as EGFR, ERBB2, FGFR, PDGF, IGF1R, and NGF. MiR-192-5p has a context dependent function, acting as a oncomiR or tumor suppressor ^**[#36 - Kim et al., 2016 | #37 - Puppo et al., 2016]**^ and in this study, the miRNA had a tumor suppressor role. NP69 cells treated with the miR-192-5p mimic showed a reduction of 40% on the transcriptional levels of *RAB2A*, along with a 43% in the migration capabilities of cells.

In this work, the *in silico* analysis of *RAB2A* showed a predicted involvement in cell cycle and cellular transcription. The RAB2A protein is a small GTPase, member of the RAS family, that controls canonical Golgi-to-Plasma membrane trafficking of a metalloprotease, MT1-MMP, essential to the acquisition of mesenchymal traits ^**[#38 - Kajiho et al., 2017, #39 - 2016]**^. *RAB2A* was previously described as a miR-192-5p target in a colon cancer model. In that context, when inhibited by miR-192-5p, *RAB2A* reduction led to inhibition of cell proliferation, migration, and invasion ^**[#40 - Zheng et al., 2019]**^.

Considering that *RAB2A* is known to induce the acquisition of mesenchymal traits, the results reported here strengthen the idea that miR-192-5p may behave as a tumor suppressor. Accordingly, this miRNA was found downregulated in several cancers; also, thyroid papillary carcinoma cells were sensitized to apoptosis and showed reduced proliferation and migration rates upon upregulation of miR-192-5p ^**[#41 - Fu et al., 2021]**^. Additionally, miR-192-5p inhibited proliferation and induced apoptosis in cervical cancer cells by targeting TRPM7, while its silencing promoted progression in endometrial cancer ^**[#42 - Ni et al., 2021]**^. In serum samples, from NPC patients, miR-192-5p was increased and proposed as a potential biomarker ^**[#43 - Zou et al., 2020]**^, however, the importance of *RAB2A* in NPC is still unpublished.

The pathway enrichment analysis performed in this study indicated that miR-192-5p regulates the TGF-β pathway (Figure 1B). The TGF-β signaling induces EMT and reduces *KHSRP* transcript, which favors the epithelial phenotype by inhibiting EMT-associated factors (Zeb1, Sai1, and Fn1, for instance), and induces miR-192-5p maturation ^**[#37 - Puppo et al., 2016]**^. Nonetheless, miR-192-5p may also inhibit TGF-β signaling via another target, Fbln2 ^**[#44 - Tang et al., 2019]**^. This miRNA was also predicted to regulate the expression of fibroblast growth factor receptor (FGF) receptors 1, 2, 2b, 3, and 4. The expression of FGFR4 was previously described to be increased in NPC tissues and cell lines, as well as correlated with higher stages and poor prognosis for NPC patients.

Worth noting, a fusion transcript from genes *FGFR3* and the gene encoding the transforming acidic coiled-coil-containing protein 3 (*TACC3*) appears to be recurrent in NPC and is related to promoting cell transformation, proliferation, and colony formation ^**[#45 - Yuan et al., 2014 p. 3]**^. The *FGFR3-TACC3* fusion also leads to overactivation of the MAPK/ERK signaling pathway, acting as an oncogenic driver, and inhibition of FGFR or MEK with BGJ398 or trametinib, respectively, blocks the transforming effects ^**[#46 - Nelson et al., 2018 | #47 - Tamura et al., 2018]**^. MiR-192-5p has been shown to target *RAB2A* in glioblastoma cells. The long non-coding RNA SOX-OT sequesters miR-192-5p, thereby increasing *RAB2A* expression and promoting glioblastoma cell growth ^**[#48 - Wang et al., 2021]**^. The expression of *RAB2A*, which is targeted by miR-192-5p, regulate the ERK1/2 pathway, sustaining its activation and increasing the nuclear accumulation of β-catenin, an effect reported to be essential to enrich CSCs in a breast cancer model ^**[#49 - Luo et al., 2015]**^.

The pathway enrichment analysis of miR-574-3p indicates a predicted effect on cellular responses to external stimuli, biological development, vesicle-mediated transport and biogenesis, cellular response to hypoxia, as well as signal transduction pathways, such as TGF-β, WNT, HIF, and TKR signaling. NP69 cells transfected with miR-574-3p mimic showed a 48% reduction on mRNA levels of *STC1*, a gene previously described to promote cancer development. In gastric cancer cells, *STC1* was increased in hypoxic, but not normoxic conditions, and when upregulated induced cell proliferation and chemoresistance^**[#50 - Wang et al., 2019]**^. This gene was also included in a proposed hypoxia signature to predict the patient outcome with bladder cancer ^**[#51 - Zhang et al., 2021]**^.

The relationship of miR-574-3p and hypoxia was previously described in a histiocytic lymphoma cell line (U937 cells) context. In a hypoxic environment, miR-574-3p is sequestered by the heterogeneous nuclear ribonucleoprotein L (hnRNP L), allowing VEGFA expression and expression of another miR-574-3p target, *EP300*, which is a transcriptional co-activator of NF-κB and HIF-1α ^**[#52 - Yao et al., 2017]**^. In a gastric cancer context, this miRNA was described as an oncomiR, increasing cell proliferation, migration, invasion, and EMT, in addition to targeting *CUL2* (another tested target), which suppresses HIF-1α ^**[#53-Ji et al., 2021]**^, which was also described to be a transcriptional regulator of *STC1* ^**[#54 - Waclawiczek et al., 2020]**^. Additionally, in gastric cancer cells, HIF-1α has been shown to directly induce the transcription of miR-574-3p, which in turn enhances HIF-1α expression by targeting CUL2. Consequently, miR-574-3p modulates the HIF/VEGF signaling pathway, thereby promoting angiogenesis under hypoxic conditions ^**[#55 - Zhang et al., 2023]**^.

## Conclusion

In conclusion, this study shows that the expression miR-192-5p (mimic) in immortalized nasopharyngeal cells inhibits the cell migration *in vitro* and the expression of *RAB2A*, a member of the RAS oncogenes. These activities suggest that this miRNA may behave as a tumor suppressor, warranting further investigation *in vivo* to assess its value in the pathogenesis of NPC.

## Acknowledgments

The authors are indebted with Prof. George Sai-Wah Tsao (Hong Kong University, China) for providing NP460hTert cells, Professor Nancy Raab-Traub (University of North Carolina, USA), for providing NP69^SV40T^ cells via Cesarman’s laboratory, and Professor Ethel Cesarman (Weill Cornell University, USA), for the insightful and unvaluable discussions about this study.

## Conflicts of Interest

The authors declare no conflict of interest.

## Ethics committee

This study has an approved Certificate of Ethical Appreciation Presentation (CAAE) #82316118.0.0000.5411 (UNESP, Botucatu School of Medicine, Committee of Research Ethics Proc. #2499053).

## Notes

**Data Availability Statement:** The authors confirm that the data supporting the findings of this study are available within the article and its supplementary materials. All material regarding this research is public available on the OpenScience Framework (OSF) platform at https://osf.io/n8xmb/

### Competing Interest Statement

The authors have declared no competing interest.

