## Supplementary Material for "Effects of human microRNAs 100-5p, 192-5p, and 574-3p on proliferation, migration, and gene expression of human immortalized nasopharyngeal cells"

**Supplementary Table S1** – Top 1% predicted target genes for miR-100-5p, miR-192-5p and miR-574-3p obtained from mirDIP platform with respective gene symbol, Uniprot code, integrated score and number of platforms that had the same prediction (number of sources).

| Gene Symbol | Uniprot | Integrated Score | Number of Sources |
|---|---|---|---|
| **miR-100-5p** | | | |
| SMARCA5 | O60264 | 0,944 | 20 |
| CTDSPL | O15194 | 0,938 | 18 |
| MTOR | P42345 | 0,932 | 20 |
| KBTBD8 | Q8NFY9 | 0,916 | 18 |
| FGFR3 | P22607 | 0,908 | 20 |
| MTMR3 | Q13615 | 0,897 | 18 |
| FZD8 | Q9H461 | 0,897 | 18 |
| HS3ST2 | Q9Y278 | 0,880 | 16 |
| MBNL1 | Q9NR56 | 0,878 | 18 |
| AGO2. | Q9UKV8 | 0,878 | 18 |
| HOXA1 | P49639 | 0,876 | 18 |
| TRIB2 | Q92519 | 0,876 | 18 |
| HS3ST3B1 | Q9Y662 | 0,852 | 15 |
| SLC44A1 | Q8WWI5 | 0,847 | 17 |
| ADCY1 | Q08828 | 0,820 | 14 |
| BAZ2A | Q9UIF9 | 0,801 | 17 |
| AP1AR | Q63HQ0 | 0,795 | 17 |
| ZZEF1 | O43149 | 0,785 | 16 |
| RAVER2 | Q9HCJ3 | 0,777 | 18 |
| INSM1 | Q01101 | 0,765 | 16 |
| PPP3CA | Q08209 | 0,740 | 16 |
| THAP2 | Q9H0W7 | 0,706 | 17 |
| IGF1R | P08069 | 0,697 | 15 |
| VLDLR | P98155 | 0,681 | 12 |
| TTC39A | Q5SRH9 | 0,674 | 15 |
| BMPR2 | Q13873 | 0,664 | 10 |
| PPP1CB | P62140 | 0,644 | 13 |
| FKBP5 | Q13451 | 0,642 | 12 |
| CYP26B1 | Q9NR63 | 0,640 | 12 |
| ST5 | P78524 | 0,637 | 15 |
| ICMT | O60725 | 0,637 | 15 |
| LRRC8B | Q6P9F7 | 0,633 | 14 |
| CLDN11 | O75508 | 0,632 | 14 |
| EPDR1 | Q9UM22 | 0,625 | 13 |
| FZD5 | Q13467 | 0,605 | 12 |
| TRIB1 | Q96RU8 | 0,589 | 11 |
| CDC25A | P30304 | 0,589 | 17 |
| PHOX2B | Q99453 | 0,588 | 8 |
| DESI2 | Q9BSY9 | 0,575 | 11 |
| FAM126B | Q8IXS8 | 0,549 | 7 |
| EPC2 | Q52LR7 | 0,546 | 7 |
| GRHL1 | Q9NZI5 | 0,538 | 12 |
| ZBTB7A | O95365 | 0,535 | 7 |
| FOXA1 | P55317 | 0,533 | 8 |
| CEP85 | Q6P2H3 | 0,518 | 9 |
| RRN3 | Q9NYV6 | 0,509 | 16 |
| NXF1 | Q9UBU9 | 0,506 | 9 |
| OGT | O15294 | 0,500 | 5 |
| RASGRP3 | Q8IV61 | 0,499 | 12 |
| TMEM30A | Q9NV96 | 0,497 | 10 |
| GMPS | P49915 | 0,493 | 8 |
| TAOK1 | Q7L7X3 | 0,492 | 10 |
| KDM6B | O15054 | 0,486 | 13 |
| RRAGD | Q9NQL2 | 0,484 | 11 |
| ATP11C | Q8NB49 | 0,484 | 13 |
| SLC14A1 | Q13336 | 0,482 | 17 |

| Gene Symbol | Uniprot | Integrated Score | Number of Sources |
|---|---|---|---|
| ST6GALNAC4 | Q9H4F1 | 0,482 | 10 |
| PI15 | O43692 | 0,482 | 9 |
| RMND5A | Q9H871 | 0,477 | 9 |
| PRDM1 | O75626 | 0,477 | 7 |
| NR6A1 | Q15406 | 0,476 | 8 |
| TARDBP | Q13148 | 0,474 | 12 |
| IREB2 | P48200 | 0,474 | 13 |
| TMPRSS13 | Q9BYE2 | 0,465 | 13 |
| C5orf22 | Q49AR2 | 0,447 | 14 |
| ADGRE2 | Q9UHX3 | 0,441 | 16 |
| IMPDH1 | P20839 | 0,436 | 15 |
| ETV3 | P41162 | 0,434 | 8 |
| RAP1B | P61224 | 0,415 | 12 |
| HES7 | Q9BYE0 | 0,414 | 12 |
| CLDN4 | O14493 | 0,414 | 7 |
| PCSK9 | Q8NBP7 | 0,406 | 12 |
| NOX4 | Q9NPH5 | 0,403 | 12 |
| ZNF845 | Q96IR2 | 0,397 | 14 |
| NTRK3 | Q16288 | 0,393 | 7 |
| PLPPR4 | Q7Z2D5 | 0,392 | 9 |
| TMEM135 | Q86UB9 | 0,392 | 8 |
| SMAD7 | O15105 | 0,391 | 9 |
| NIPBL | Q6KC79 | 0,390 | 6 |
| ANKRD28 | O15084 | 0,389 | 13 |
| RNF144B | Q7Z419 | 0,386 | 11 |
| ZNF197 | O14709 | 0,386 | 12 |
| **miR-192-5p** | | | |
| PKP4 | Q99569 | 0.890 | 19 |
| H3F3B | P84243 | 0.883 | 19 |
| PABPC4 | Q13310 | 0.875 | 20 |
| RAB2A | P61019 | 0.863 | 21 |
| ARFGEF1 | Q9Y6D6 | 0.858 | 18 |
| EREG | O14944 | 0.842 | 20 |
| MSN | P26038 | 0.835 | 17 |
| ALCAM | Q13740 | 0.832 | 18 |
| DYRK1A | Q13627 | 0.824 | 20 |
| FRMD4B | Q9Y2L6 | 0.821 | 19 |
| LPAR4 | Q99677 | 0.820 | 18 |
| WDR44 | Q5JSH3 | 0.817 | 16 |
| CCNT2 | O60583 | 0.812 | 19 |
| DBT | P11182 | 0.811 | 20 |
| GPR22 | Q99680 | 0.810 | 16 |
| FNDC3B | Q53EP0 | 0.809 | 19 |
| BHLHE22 | Q8NFJ8 | 0.803 | 17 |
| CTNNBIP1 | Q9NSA3 | 0.801 | 21 |

| Gene Symbol | Uniprot | Integrated Score | Number of Sources |
|---|---|---|---|
| RB1 | P06400 | 0.801 | 18 |
| NIPBL | Q6KC79 | 0.799 | 19 |
| ZNF536 | O15090 | 0.787 | 17 |
| MIPOL1 | Q8TD10 | 0.787 | 17 |
| MTMR4 | Q9NYA4 | 0.785 | 18 |
| PRKAR1A | P10644 | 0.771 | 18 |
| DDX6 | P26196 | 0.770 | 19 |
| DICER1 | Q9UPY3 | 0.767 | 18 |
| PPP1R3D | O95685 | 0.766 | 17 |
| RUNX1T1 | Q06455 | 0.766 | 15 |
| ZEB2 | O60315 | 0.765 | 18 |
| SRSF6 | Q13247 | 0.758 | 17 |
| DYRK3 | O43781 | 0.756 | 18 |
| WNK1 | Q9H4A3 | 0.755 | 17 |
| ZBTB34 | Q8NCN2 | 0.752 | 16 |
| TDG | Q13569 | 0.748 | 16 |
| CBL | P22681 | 0.746 | 17 |
| CTCF | P49711 | 0.742 | 19 |
| ADGRL3 | Q9HAR2 | 0.739 | 17 |
| FABP3 | P05413 | 0.718 | 13 |
| BLCAP | P62952 | 0.712 | 17 |
| NKAIN2 | Q5VXU1 | 0.709 | 15 |
| PDP1 | Q9P0J1 | 0.704 | 16 |
| SLC39A6 | Q13433 | 0.701 | 15 |
| NAA50 | Q9GZZ1 | 0.699 | 18 |
| OLIG3 | Q7RTU3 | 0.698 | 14 |
| SLC19A2 | O60779 | 0.694 | 19 |
| NRIP1 | P48552 | 0.691 | 13 |
| ALX1 | Q15699 | 0.691 | 11 |
| SOAT1 | P35610 | 0.689 | 16 |
| NCOA3 | Q9Y6Q9 | 0.685 | 16 |
| SLC5A3 | P53794 | 0.685 | 14 |
| KPNA6 | O60684 | 0.683 | 16 |
| FAM229B | Q4G0N7 | 0.679 | 14 |
| ZNF280C | Q8ND82 | 0.678 | 15 |
| ACVR2A | P27037 | 0.675 | 17 |
| IKZF2 | Q9UKS7 | 0.673 | 19 |
| RUNX1 | Q01196 | 0.671 | 16 |
| CHD7 | Q9P2D1 | 0.668 | 13 |
| DDX50 | Q9BQ39 | 0.667 | 14 |
| B3GALNT1 | O75752 | 0.665 | 19 |
| SLMAP | Q14BN4 | 0.663 | 17 |
| TRERF1 | Q96PN7 | 0.662 | 15 |
| CREB5 | Q02930 | 0.654 | 16 |
| RPAP2 | Q8IXW5 | 0.653 | 15 |

| Gene Symbol | Uniprot | Integrated Score | Number of Sources |
|---|---|---|---|
| ACVR2B | Q13705 | 0.653 | 15 |
| ARL4C | P56559 | 0.652 | 14 |
| COL5A1 | P20908 | 0.652 | 15 |
| C6orf106 | Q9H6K1 | 0.649 | 16 |
| DDX3X | O00571 | 0.649 | 12 |
| CLSTN1 | O94985 | 0.649 | 16 |
| PRKACB | P22694 | 0.648 | 15 |
| TRIM44 | Q96DX7 | 0.647 | 14 |
| ZC3HAV1 | Q7Z2W4 | 0.646 | 18 |
| BRD3 | Q15059 | 0.645 | 17 |
| KCNA7 | Q96RP8 | 0.645 | 17 |
| PTPRT | O14522 | 0.643 | 15 |
| ZFHX4 | Q86UP3 | 0.641 | 11 |
| KCNK1 | O00180 | 0.640 | 15 |
| PHTF2 | Q8N3S3 | 0.640 | 13 |
| LARP4 | Q71RC2 | 0.638 | 18 |
| ARHGAP36 | Q6ZRI8 | 0.636 | 15 |
| CBLN4 | Q9NTU7 | 0.636 | 16 |
| GAPVD1 | Q14C86 | 0.633 | 16 |
| SERTAD2 | Q14140 | 0.630 | 9 |
| LRRFIP1 | Q32MZ4 | 0.630 | 15 |
| MED14 | O60244 | 0.629 | 15 |
| ADGRG6 | Q86SQ4 | 0.628 | 13 |
| ANKRD44 | Q8N8A2 | 0.627 | 13 |
| TMTC3 | Q6ZXV5 | 0.624 | 14 |
| ATF1 | P18846 | 0.623 | 12 |
| SYT6 | Q5T7P8 | 0.623 | 16 |
| ZBTB4 | Q9P1Z0 | 0.623 | 14 |
| PCDH9 | Q9HC56 | 0.622 | 14 |
| SCN3A | Q9NY46 | 0.620 | 16 |
| XIAP | P98170 | 0.619 | 13 |
| AP3M2 | P53677 | 0.618 | 16 |
| CDC7 | O00311 | 0.617 | 17 |
| CNGB3 | Q9NQW8 | 0.616 | 16 |
| KHDRBS3 | O75525 | 0.613 | 15 |
| KIF1B | O60333 | 0.612 | 17 |
| TYMS | P04818 | 0.612 | 15 |
| HIGD1A | Q9Y241 | 0.612 | 14 |
| RICTOR | Q6R327 | 0.610 | 14 |
| PRKD3 | O94806 | 0.610 | 15 |
| CPEB4 | Q17RY0 | 0.608 | 9 |
| EFNB2 | P52799 | 0.608 | 13 |
| ZMAT3 | Q9HA38 | 0.605 | 16 |
| MFAP3 | P55082 | 0.605 | 14 |
| ARHGEF39 | Q8N4T4 | 0.604 | 12 |

| Gene Symbol | Uniprot | Integrated Score | Number of Sources |
|---|---|---|---|
| EMC7 | Q9NPA0 | 0.599 | 11 |
| LIMS1 | P48059 | 0.599 | 15 |
| ANKS1A | Q92625 | 0.598 | 16 |
| C4orf46 | Q504U0 | 0.597 | 13 |
| ARHGAP19 | Q14CB8 | 0.597 | 15 |
| RNF141 | Q8WVD5 | 0.595 | 15 |
| ALKBH8 | Q96BT7 | 0.595 | 15 |
| L2HGDH | Q9H9P8 | 0.594 | 17 |
| SEMA4D | Q92854 | 0.591 | 15 |
| TMEM106B | Q9NUM4 | 0.591 | 16 |
| IER5 | Q5VY09 | 0.591 | 13 |
| RABGAP1 | Q9Y3P9 | 0.590 | 13 |
| ENC1 | O14682 | 0.585 | 11 |
| TRIP13 | Q15645 | 0.585 | 14 |
| CXCL2 | P19875 | 0.584 | 17 |
| C1orf21 | Q9H246 | 0.583 | 12 |
| USP1 | O94782 | 0.582 | 12 |
| TMEM30A | Q9NV96 | 0.582 | 14 |
| CCDC121 | Q6ZUS5 | 0.580 | 14 |
| CCDC152 | Q4G0S7 | 0.579 | 13 |
| SMARCAD1 | Q9H4L7 | 0.578 | 13 |
| AMER1 | Q5JTC6 | 0.578 | 15 |
| FGD5 | Q6ZNL6 | 0.577 | 15 |
| SH3RF3 | Q8TEJ3 | 0.576 | 13 |
| CDON | Q4KMG0 | 0.575 | 13 |
| EGR1 | P18146 | 0.575 | 15 |
| RIC8B | Q9NVN3 | 0.574 | 14 |
| TOR1AIP1 | Q5JTV8 | 0.573 | 16 |
| APLN | Q9ULZ1 | 0.573 | 14 |
| ZNF451 | Q9Y4E5 | 0.573 | 12 |
| FAM129A | Q9BZQ8 | 0.572 | 16 |
| SLC11A2 | P49281 | 0.571 | 14 |
| MYLK | Q15746 | 0.570 | 14 |
| WASHC4 | Q2M389 | 0.569 | 15 |
| CEP70 | Q8NHQ1 | 0.568 | 15 |
| UBL3 | O95164 | 0.566 | 13 |
| IGDCC3 | Q8IVU1 | 0.565 | 9 |
| NOD2 | Q9HC29 | 0.563 | 14 |
| SMC5 | Q8IY18 | 0.561 | 11 |
| HIF1AN | Q9NWT6 | 0.560 | 15 |
| ARL2BP | Q9Y2Y0 | 0.559 | 14 |
| DCC | P43146 | 0.557 | 14 |
| GABPB1 | Q06547 | 0.556 | 13 |
| HIBADH | P31937 | 0.554 | 14 |
| TSHZ2 | Q9NRE2 | 0.554 | 14 |

| Gene Symbol | Uniprot | Integrated Score | Number of Sources |
|---|---|---|---|
| NSD2 | O96028 | 0.552 | 14 |
| KIF5B | P33176 | 0.551 | 12 |
| LPAR1 | Q92633 | 0.550 | 15 |
| SAMD4A | Q9UPU9 | 0.549 | 16 |
| PLXNB2 | O15031 | 0.548 | 13 |
| CERS6 | Q6ZMG9 | 0.548 | 11 |
| DIXDC1 | Q155Q3 | 0.548 | 14 |
| SPIN1 | Q9Y657 | 0.547 | 14 |
| DCUN1D4 | Q92564 | 0.546 | 15 |
| FAM167A | Q96KS9 | 0.546 | 13 |
| GRHL1 | Q9NZI5 | 0.546 | 13 |
| PCDH19 | Q8TAB3 | 0.545 | 14 |
| CUL3 | Q13618 | 0.545 | 14 |
| F13A1 | P00488 | 0.543 | 15 |
| RFWD3 | Q6PCD5 | 0.543 | 14 |
| ZFHX3 | Q15911 | 0.542 | 11 |
| FOXN1 | O15353 | 0.542 | 9 |
| NFAT5 | O94916 | 0.541 | 13 |
| FGF7 | P21781 | 0.539 | 15 |
| SPARC | P09486 | 0.538 | 12 |
| SCN1A | P35498 | 0.535 | 12 |
| GLP1R | P43220 | 0.534 | 10 |
| ZPBP2 | Q6X784 | 0.534 | 10 |
| RAP1GAP2 | Q684P5 | 0.534 | 9 |
| SRSF3 | P84103 | 0.534 | 11 |
| CSMD3 | Q7Z407 | 0.532 | 14 |
| KMT2A | Q03164 | 0.532 | 14 |
| KIF20B | Q96Q89 | 0.531 | 15 |
| PERP | Q96FX8 | 0.531 | 14 |
| DLG5 | Q8TDM6 | 0.530 | 8 |
| DIAPH1 | O60610 | 0.528 | 14 |
| WWC2 | Q6AWC2 | 0.528 | 11 |
| CNOT6L | Q96LI5 | 0.528 | 15 |
| ABI2 | Q9NYB9 | 0.528 | 16 |
| NIN | Q8N4C6 | 0.527 | 14 |
| RAD54B | Q9Y620 | 0.526 | 11 |
| ARMC8 | Q8IUR7 | 0.525 | 15 |
| APPBP2 | Q92624 | 0.524 | 12 |
| KLHL15 | Q96M94 | 0.523 | 12 |
| SRGAP3 | O43295 | 0.522 | 10 |
| KCNQ5 | Q9NR82 | 0.522 | 15 |
| ASXL2 | Q76L83 | 0.522 | 13 |
| SLC30A9 | Q6PML9 | 0.521 | 12 |
| CRK | P46108 | 0.521 | 14 |
| C10orf90 | Q96M02 | 0.520 | 14 |
| ARIH1 | Q9Y4X5 | 0.519 | 14 |
| FAM234B | A2RU67 | 0.519 | 13 |
| CD164 | Q04900 | 0.519 | 15 |
| NAV1 | Q8NEY1 | 0.518 | 12 |
| SNRPG | P62308 | 0.518 | 14 |
| CRX | O43186 | 0.518 | 17 |
| NSF | P46459 | 0.517 | 14 |
| WSCD2 | Q2TBF2 | 0.515 | 12 |
| VAPB | O95292 | 0.515 | 12 |
| SEMA3A | Q14563 | 0.513 | 12 |
| KIDINS220 | Q9ULH0 | 0.513 | 14 |
| CALD1 | Q05682 | 0.513 | 14 |
| IL6ST | P40189 | 0.513 | 10 |
| MAP3K1 | Q13233 | 0.513 | 13 |
| SREK1IP1 | Q8N9Q2 | 0.513 | 15 |
| ITGAV | P06756 | 0.511 | 15 |
| RSAD2 | Q8WXG1 | 0.511 | 14 |
| CUL5 | Q93034 | 0.510 | 15 |
| ASB6 | Q9NWX5 | 0.510 | 11 |
| RPRD1B | Q9NQG5 | 0.510 | 13 |
| C8orf46 | Q8TAG6 | 0.509 | 16 |
| KLHL42 | Q9P2K6 | 0.509 | 14 |
| FAM98A | Q8NCA5 | 0.509 | 14 |
| ATAD2B | Q9ULI0 | 0.507 | 13 |
| ABCG5 | Q9H222 | 0.506 | 14 |
| DYNC2H1 | Q8NCM8 | 0.506 | 14 |
| ZNF654 | Q8IZM8 | 0.505 | 13 |
| CAV1 | Q03135 | 0.504 | 14 |
| SEC63 | Q9UGP8 | 0.503 | 13 |
| KPNA4 | O00629 | 0.503 | 10 |
| CRTC2 | Q53ET0 | 0.503 | 9 |
| MGEA5 | O60502 | 0.502 | 13 |
| GCLM | P48507 | 0.502 | 13 |
| DIAPH2 | O60879 | 0.501 | 9 |
| ACPP | P15309 | 0.501 | 14 |
| ACTBL2 | Q562R1 | 0.500 | 12 |
| PTBP2 | Q9UKA9 | 0.499 | 11 |
| NTRK2 | Q16620 | 0.498 | 13 |
| GAD1 | Q99259 | 0.497 | 14 |
| CRISP1 | P54107 | 0.496 | 14 |
| CADM1 | Q9BY67 | 0.496 | 14 |
| IGDCC4 | Q8TDY8 | 0.494 | 9 |
| HOXA10 | P31260 | 0.494 | 9 |
| ELOA | Q14241 | 0.493 | 9 |
| UMODL1 | Q5DID0 | 0.493 | 15 |

| Gene Symbol | Uniprot | Integrated Score | Number of Sources |
|---|---|---|---|
| DPP10 | Q8N608 | 0.493 | 11 |
| MAPK1 | P28482 | 0.492 | 11 |
| DPYSL2 | Q16555 | 0.490 | 13 |
| FAM46A | Q96IP4 | 0.490 | 15 |
| CYCS | P99999 | 0.490 | 15 |
| CACNA1I | Q9P0X4 | 0.489 | 12 |
| TCF7 | P36402 | 0.489 | 16 |
| CLIP3 | Q96DZ5 | 0.488 | 14 |
| GABBR2 | O75899 | 0.488 | 10 |
| DIS3L | Q8TF46 | 0.487 | 13 |
| NEFL | P07196 | 0.487 | 11 |
| GOLGA8A | A7E2F4 | 0.486 | 13 |
| ZBTB38 | Q8NAP3 | 0.486 | 14 |
| PARP8 | Q8N3A8 | 0.486 | 12 |
| INO80D | Q53TQ3 | 0.486 | 14 |
| IDS | P22304 | 0.486 | 14 |
| TRPM7 | Q96QT4 | 0.486 | 10 |
| TPM1 | P09493 | 0.486 | 12 |
| STAG1 | Q8WVM7 | 0.484 | 10 |
| MACF1 | Q9UPN3 | 0.483 | 10 |
| NRIP3 | Q9NQ35 | 0.483 | 15 |
| PCGF5 | Q86SE9 | 0.483 | 14 |
| DCAF4L2 | Q8NA75 | 0.482 | 13 |
| CAMTA1 | Q9Y6Y1 | 0.481 | 11 |
| MEF2C | Q06413 | 0.481 | 11 |
| SLC24A4 | Q8NFF2 | 0.481 | 14 |
| SOCS6 | O14544 | 0.480 | 12 |
| CAMSAP2 | Q08AD1 | 0.480 | 12 |
| PPP1CB | P62140 | 0.480 | 14 |
| MCM10 | Q7L590 | 0.479 | 14 |
| LYRM7 | Q5U5X0 | 0.478 | 13 |
| ZMYM1 | Q5SVZ6 | 0.478 | 13 |
| BCAP29 | Q9UHQ4 | 0.478 | 14 |
| CTH | P32929 | 0.477 | 11 |
| TCTEX1D1 | Q8N7M0 | 0.477 | 10 |
| INPP4A | Q96PE3 | 0.477 | 14 |
| CCDC47 | Q96A33 | 0.476 | 14 |
| DNM3 | Q9UQ16 | 0.476 | 14 |
| SH2D1A | O60880 | 0.476 | 16 |
| SH3TC2 | Q8TF17 | 0.476 | 12 |
| SC5D | O75845 | 0.476 | 14 |
| MECP2 | P51608 | 0.475 | 12 |
| ESR1 | P03372 | 0.475 | 14 |
| FZD4 | Q9ULV1 | 0.474 | 14 |
| SYNPO2 | Q9UMS6 | 0.474 | 13 |

| Gene Symbol | Uniprot | Integrated Score | Number of Sources |
|---|---|---|---|
| RETREG2 | Q8NC44 | 0.473 | 13 |
| DIEXF | Q68CQ4 | 0.473 | 12 |
| NID1 | P14543 | 0.472 | 13 |
| TRIM2 | Q9C040 | 0.471 | 12 |
| GMEB1 | Q9Y692 | 0.471 | 11 |
| NR6A1 | Q15406 | 0.470 | 10 |
| NEK1 | Q96PY6 | 0.470 | 13 |
| TRAF4 | Q9BUZ4 | 0.469 | 7 |
| HBS1L | Q9Y450 | 0.467 | 16 |
| ZNF136 | P52737 | 0.467 | 13 |
| ATP2C1 | P98194 | 0.467 | 14 |
| NUDT15 | Q9NV35 | 0.467 | 13 |
| BCL2L11 | O43521 | 0.465 | 14 |
| CTTNBP2NL | Q9P2B4 | 0.464 | 13 |
| PTPRE | P23469 | 0.464 | 10 |
| BRWD1 | Q9NSI6 | 0.463 | 13 |
| PNPT1 | Q8TCS8 | 0.463 | 15 |
| PCDH17 | O14917 | 0.462 | 8 |
| CCDC171 | Q6TFL3 | 0.462 | 9 |
| BMPR2 | Q13873 | 0.461 | 12 |
| TRAF5 | O00463 | 0.461 | 13 |
| DCK | P27707 | 0.461 | 7 |
| BMPER | Q8N8U9 | 0.460 | 15 |
| GABPB2 | Q8TAK5 | 0.460 | 11 |
| TMEM67 | Q5HYA8 | 0.460 | 15 |
| ATF3 | P18847 | 0.460 | 14 |
| IL1RAP | Q9NPH3 | 0.460 | 14 |
| AFF2 | P51816 | 0.459 | 13 |
| DNAH5 | Q8TE73 | 0.459 | 11 |
| PRNP | F7VJQ1 | 0.459 | 12 |
| KCNK3 | O14649 | 0.459 | 11 |
| MYO6 | Q9UM54 | 0.458 | 14 |
| FAM199X | Q6PEV8 | 0.457 | 12 |
| FZD9 | O00144 | 0.457 | 7 |
| SIK1 | P57059 | 0.456 | 12 |
| PPP2CB | P62714 | 0.455 | 14 |
| PMP2 | P02689 | 0.455 | 14 |
| RNF6 | Q9Y252 | 0.454 | 9 |
| TRAIP | Q9BWF2 | 0.454 | 13 |
| CCND2 | P30279 | 0.454 | 13 |
| TBC1D22B | Q9NU19 | 0.454 | 13 |
| LARP1B | Q659C4 | 0.453 | 12 |
| DLGAP1 | O14490 | 0.453 | 14 |
| RAD1 | O60671 | 0.453 | 14 |
| OSBPL10 | Q9BXB5 | 0.452 | 10 |

| Gene Symbol | Uniprot | Integrated Score | Number of Sources |
|---|---|---|---|
| NKX2-5 | P52952 | 0.452 | 12 |
| LOXL2 | Q9Y4K0 | 0.451 | 14 |
| C2orf71 | A6NGG8 | 0.451 | 14 |
| PSMD5 | Q16401 | 0.451 | 12 |
| PRKG1 | Q13976 | 0.449 | 9 |
| C1D | Q13901 | 0.448 | 13 |
| MUM1L1 | Q5H9M0 | 0.447 | 12 |
| MKL2 | Q9ULH7 | 0.447 | 12 |
| INAVA | Q3KP66 | 0.447 | 14 |
| LRCH2 | Q5VUJ6 | 0.446 | 11 |
| C16orf46 | Q6P387 | 0.446 | 14 |
| SLC35F3 | Q8IY50 | 0.446 | 13 |
| MIS12 | Q9H081 | 0.446 | 13 |
| OTUD3 | Q5T2D3 | 0.446 | 12 |
| CRNKL1 | Q9BZJ0 | 0.446 | 14 |
| NAB1 | Q13506 | 0.446 | 10 |
| ATXN7 | O15265 | 0.445 | 15 |
| GOLGA8B | A8MQT2 | 0.445 | 11 |
| IGF1 | P05019 | 0.445 | 12 |
| SGCD | Q92629 | 0.445 | 12 |
| BCAT1 | P54687 | 0.444 | 13 |
| ATP8B4 | Q8TF62 | 0.444 | 13 |
| COPS7A | Q9UBW8 | 0.444 | 14 |
| NKX3-1 | Q99801 | 0.444 | 13 |
| CXCR5 | P32302 | 0.443 | 9 |
| RALB | P11234 | 0.443 | 12 |
| PHACTR2 | O75167 | 0.443 | 13 |
| ICK | Q9UPZ9 | 0.443 | 12 |
| CRYBG3 | Q68DQ2 | 0.442 | 11 |
| CENPBD1 | B2RD01 | 0.442 | 12 |
| REPS2 | Q8NFH8 | 0.441 | 12 |
| ORC4 | O43929 | 0.439 | 14 |
| PRKCQ | Q04759 | 0.439 | 13 |
| SNTB2 | Q13425 | 0.438 | 13 |
| MED28 | Q9H204 | 0.438 | 15 |
| CSTF1 | Q05048 | 0.437 | 14 |
| SLAMF7 | Q9NQ25 | 0.437 | 13 |
| ONECUT2 | O95948 | 0.436 | 13 |
| GNG3 | P63215 | 0.436 | 13 |
| STX7 | O15400 | 0.436 | 12 |
| VPS37B | Q9H9H4 | 0.436 | 14 |
| UBA6 | A0AVT1 | 0.435 | 14 |
| JCHAIN | P01591 | 0.435 | 12 |
| WFDC13 | Q8IUB5 | 0.434 | 13 |
| C14orf105 | Q9NVL8 | 0.434 | 11 |
| RFX6 | Q8HWS3 | 0.434 | 8 |
| NKRF | O15226 | 0.433 | 11 |
| SYNJ2 | O15056 | 0.433 | 12 |
| JAGN1 | Q8N5M9 | 0.433 | 14 |
| LEFTY2 | O00292 | 0.432 | 15 |
| GALNTL6 | Q49A17 | 0.432 | 7 |
| ACADSB | P45954 | 0.432 | 12 |
| EIF5A2 | Q9GZV4 | 0.432 | 8 |
| MAP1LC3B | Q9GZQ8 | 0.432 | 14 |
| VCAN | P13611 | 0.431 | 12 |
| ANAPC16 | Q96DE5 | 0.431 | 11 |
| ZDHHC2 | Q9UIJ5 | 0.431 | 10 |
| BIN2 | Q9UBW5 | 0.431 | 13 |
| TRIM23 | P36406 | 0.430 | 13 |
| LHX6 | Q9UPM6 | 0.430 | 13 |
| SCD | O00767 | 0.430 | 12 |
| SDE2 | Q6IQ49 | 0.429 | 13 |
| PDHB | P11177 | 0.429 | 7 |
| YY1 | P25490 | 0.428 | 7 |
| USP45 | Q70EL2 | 0.428 | 13 |
| XPO4 | Q9C0E2 | 0.428 | 9 |
| POLR3F | Q9H1D9 | 0.428 | 14 |
| C3orf14 | Q9HBI5 | 0.428 | 8 |
| ZC3H12B | Q5HYM0 | 0.428 | 12 |
| MCM6 | Q14566 | 0.428 | 13 |
| NCAM1 | P13591 | 0.427 | 12 |
| RGMB | Q6NW40 | 0.427 | 13 |
| MKNK2 | Q9HBH9 | 0.427 | 15 |
| LINC01554 | Q52M75 | 0.426 | 6 |
| ADCY7 | P51828 | 0.426 | 13 |
| ABHD2 | P08910 | 0.426 | 12 |
| GLYCTK | Q8IVS8 | 0.426 | 12 |
| DCAF8 | Q5TAQ9 | 0.426 | 13 |
| PIP4K2B | P78356 | 0.426 | 10 |
| C5orf30 | Q96GV9 | 0.425 | 12 |
| ATF7 | P17544 | 0.424 | 12 |
| RCOR1 | Q9UKL0 | 0.424 | 11 |
| CWC25 | Q9NXE8 | 0.424 | 13 |
| B4GALT2 | O60909 | 0.423 | 10 |
| SOD2 | P04179 | 0.423 | 13 |
| FBXO11 | Q86XK2 | 0.423 | 9 |
| ZBTB39 | O15060 | 0.422 | 11 |
| RALGPS1 | Q5JS13 | 0.422 | 12 |
| GAS7 | O60861 | 0.422 | 12 |
| ENOX2 | Q16206 | 0.421 | 14 |

| Gene Symbol | Uniprot | Integrated Score | Number of Sources | Gene Symbol | Uniprot | Integrated Score | Number of Sources |
|---|---|---|---|---|---|---|---|
| SLC35B4 | Q969S0 | 0.421 | 12 | DNAJC24 | Q6P3W2 | 0.409 | 9 |
| TAB2 | Q9NYJ8 | 0.420 | 10 | PRRG1 | O14668 | 0.408 | 14 |
| SYAP1 | Q96A49 | 0.420 | 11 | SPCS3 | P61009 | 0.408 | 12 |
| KRAS | P01116 | 0.419 | 9 | PLEKHA1 | Q9HB21 | 0.408 | 9 |
| PCDHGB7 | Q9Y5F8 | 0.419 | 9 | PRKAA2 | P54646 | 0.407 | 14 |
| G6PC | P35575 | 0.418 | 12 | MYO9B | Q13459 | 0.407 | 12 |
| MMP16 | P51512 | 0.418 | 7 | NIPAL1 | Q6NVV3 | 0.407 | 7 |
| DYRK2 | Q92630 | 0.418 | 12 | OXR1 | Q8N573 | 0.407 | 10 |
| HOMER1 | Q86YM7 | 0.418 | 13 | HDAC2 | Q92769 | 0.407 | 12 |
| TOP1 | P11387 | 0.417 | 10 | GOLGA6A | Q9NYA3 | 0.406 | 13 |
| PPP1R18 | Q6NYC8 | 0.417 | 15 | DNAJC19 | Q96DA6 | 0.406 | 6 |
| FGF2 | P09038 | 0.416 | 10 | UBE2W | Q96B02 | 0.405 | 12 |
| SLC23A3 | Q6PIS1 | 0.416 | 8 | DLD | P09622 | 0.404 | 10 |
| STRIP2 | Q9ULQ0 | 0.416 | 11 | STXBP4 | Q6ZWJ1 | 0.404 | 12 |
| GPR137B | O60478 | 0.416 | 14 | AP1S3 | Q96PC3 | 0.404 | 12 |
| CSNK1G3 | Q9Y6M4 | 0.416 | 8 | ANGEL2 | Q5VTE6 | 0.404 | 11 |
| ATP1A2 | P50993 | 0.415 | 12 | PTGER3 | P43115 | 0.403 | 13 |
| KDM1B | Q8NB78 | 0.415 | 13 | RNF8 | O76064 | 0.402 | 13 |
| SLC16A14 | Q7RTX9 | 0.415 | 10 | ZBTB24 | O43167 | 0.402 | 10 |
| MYO5B | Q9ULV0 | 0.414 | 13 | MYPN | Q86TC9 | 0.402 | 13 |
| MYO1E | Q12965 | 0.414 | 15 | TPD52L3 | Q96J77 | 0.402 | 13 |
| ARHGEF15 | O94989 | 0.414 | 12 | PRDM10 | Q9NQV6 | 0.402 | 13 |
| VSX1 | Q9NZR4 | 0.414 | 12 | CFAP65 | Q6ZU64 | 0.401 | 13 |
| MPZ | P25189 | 0.413 | 14 | SLAIN2 | Q9P270 | 0.401 | 12 |
| PTCHD1 | Q96NR3 | 0.413 | 8 | B3GAT1 | Q9P2W7 | 0.400 | 13 |
| IFFO2 | Q5TF58 | 0.413 | 14 | CRLF3 | Q8IUI8 | 0.399 | 13 |
| PCNP | Q8WW12 | 0.413 | 9 | SNX33 | Q8WV41 | 0.399 | 9 |
| NCEH1 | Q6PIU2 | 0.412 | 13 | C11orf68 | Q9H3H3 | 0.398 | 13 |
| U2SURP | O15042 | 0.412 | 11 | FGFR1OP | O95684 | 0.398 | 9 |
| ZFYVE26 | Q68DK2 | 0.412 | 12 | TRPS1 | Q9UHF7 | 0.398 | 9 |
| EXOC4 | Q96A65 | 0.412 | 13 | SERF1A | O75920 | 0.397 | 12 |
| FAM210B | Q96KR6 | 0.412 | 12 | SRRM4 | A7MD48 | 0.397 | 12 |
| TIGAR | Q9NQ88 | 0.412 | 9 | MBTD1 | Q05BQ5 | 0.397 | 11 |
| EIF1 | P41567 | 0.411 | 7 | SHANK2 | Q9UPX8 | 0.396 | 10 |
| SLC22A15 | Q8IZD6 | 0.411 | 12 | MRAP | Q8TCY5 | 0.396 | 13 |
| PPM1A | P35813 | 0.411 | 11 | USP48 | Q86UV5 | 0.396 | 11 |
| ZC3H6 | P61129 | 0.411 | 12 | NSD1 | Q96L73 | 0.396 | 13 |
| DCX | O43602 | 0.410 | 13 | PDSS2 | Q86YH6 | 0.396 | 10 |
| BCO2 | Q9BYV7 | 0.410 | 12 | HOXB9 | P17482 | 0.395 | 12 |
| HOOK3 | Q86VS8 | 0.410 | 10 | MRVI1 | Q9Y6F6 | 0.395 | 12 |
| TRIM58 | Q8NG06 | 0.410 | 13 | ERCC3 | P19447 | 0.395 | 14 |
| TAOK1 | Q7L7X3 | 0.409 | 9 | KIAA1147 | A4D1U4 | 0.394 | 12 |
| B3GALNT2 | Q8NCR0 | 0.409 | 11 | NPAS3 | Q8IXF0 | 0.394 | 10 |
| MS4A1 | P11836 | 0.409 | 12 | CEP85L | Q5SZL2 | 0.394 | 13 |
| DPYSL3 | Q14195 | 0.409 | 14 | LVRN | Q6Q4G3 | 0.393 | 13 |

| Gene Symbol | Uniprot | Integrated Score | Number of Sources |
|---|---|---|---|
| FURIN | P09958 | 0.393 | 14 |
| KRBA2 | Q6ZNG9 | 0.393 | 8 |
| RBM26 | Q5T8P6 | 0.392 | 8 |
| UBE2QL1 | A1L167 | 0.392 | 8 |
| SLC7A6 | Q92536 | 0.392 | 11 |
| OPRM1 | P35372 | 0.391 | 12 |
| CD47 | Q08722 | 0.391 | 12 |
| PHF20 | Q9BVI0 | 0.391 | 8 |
| VCPIP1 | Q96JH7 | 0.391 | 9 |
| ATM | Q13315 | 0.391 | 10 |
| KPNA1 | P52294 | 0.390 | 10 |
| DHTKD1 | Q96HY7 | 0.390 | 12 |
| CACNA1C | Q13936 | 0.390 | 13 |
| APPL1 | Q9UKG1 | 0.390 | 11 |
| STRBP | Q96SI9 | 0.390 | 13 |
| CREBRF | Q8IUR6 | 0.390 | 9 |
| MEGF10 | Q96KG7 | 0.389 | 13 |
| SSH2 | Q76I76 | 0.389 | 11 |
| LMTK2 | Q8IWU2 | 0.389 | 7 |
| RAB6A | P20340 | 0.388 | 10 |
| CEBPG | P53567 | 0.388 | 11 |
| TUB | P50607 | 0.388 | 12 |
| PHF20L1 | A8MW92 | 0.388 | 9 |
| ARHGAP29 | Q52LW3 | 0.387 | 12 |
| CST9 | Q5W186 | 0.387 | 11 |
| SERF1B | O75920 | 0.385 | 12 |
| VPS53 | Q5VIR6 | 0.385 | 11 |
| TNRC6B | Q9UPQ9 | 0.384 | 10 |
| ASPH | Q12797 | 0.383 | 11 |
| ADAMTS3 | O15072 | 0.383 | 11 |
| ATP6V1C2 | Q8NEY4 | 0.383 | 8 |
| MYH15 | Q9Y2K3 | 0.382 | 12 |
| **miR-574-3p** | | | |
| CUL2 | Q13617 | 0,601 | 16 |
| CLTC | Q00610 | 0,484 | 13 |
| STRN3 | Q13033 | 0,476 | 13 |
| SNRK | Q9NRH2 | 0,456 | 13 |
| TMCC1 | O94876 | 0,410 | 11 |
| TP53TG3 | Q9ULZ0 | 0,405 | 11 |
| ACVR2B | Q13705 | 0,403 | 12 |
| SLC6A3 | Q01959 | 0,400 | 13 |
| STC1 | P52823 | 0,384 | 12 |
| FBXL5 | Q9UKA1 | 0,384 | 14 |

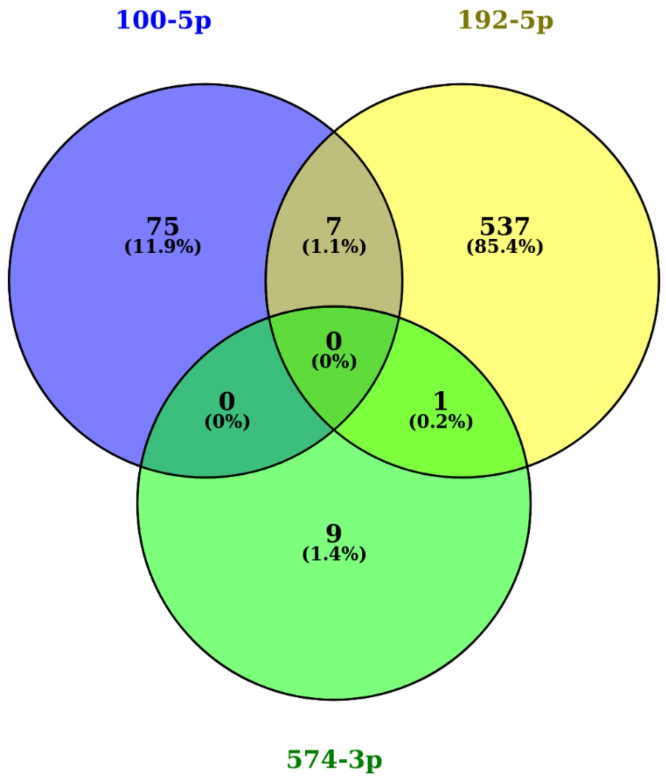


**Supplementary Figure S1** – Target prediction analysis of miR-100-5p, miR-192-5p and miR-574-3p shows that miR-100-5p (blue) has 82 targets, miR-192-5p (yellow) has 545 targets and miR-574-3p (green) has 10 targets. There is 7 targets shared between miR-100-5p and miR-192-5p, and 1 shared between miR-192-5p and miR-574-3p.

**Supplementary Table S2 –** Pathway Enrichment Analysis performed with Reactome platform using only predicted genes with the top 1% integrated scores generated in mirDIP target prediction software.

| Reactome Pathway | Ratio of Protein In Pathway | Number of Protein In Pathway | Protein from Gene Set | P-value |
|---|---|---|---|---|
| **miR-100-5p** | | | | |
| VLDLR internalisation and degradation | 0,0015 | 12 | 2 | 1,58E-03 |
| Macroautophagy | 0,0074 | 60 | 3 | 3,07E-03 |
| Choline catabolism | 0,0001 | 1 | 1 | 4,82E-03 |
| mTORC1-mediated signalling | 0,0027 | 22 | 2 | 5,14E-03 |
| Signaling by BMP | 0,0028 | 23 | 2 | 5,60E-03 |
| VLDL interactions | 0,0028 | 23 | 2 | 5,60E-03 |
| PI3K Cascade | 0,0098 | 79 | 3 | 6,57E-03 |
| HS-GAG biosynthesis | 0,0032 | 26 | 2 | 7,09E-03 |
| Downregulation of TGF-beta receptor signaling | 0,0032 | 26 | 2 | 7,09E-03 |
| IRS-related events triggered by IGF1R | 0,0314 | 254 | 5 | 7,26E-03 |
| IGF1R signaling cascade | 0,0314 | 254 | 5 | 7,26E-03 |
| Signaling by Type 1 Insulin-like Growth Factor 1 Receptor (IGF1R) | 0,0316 | 255 | 5 | 7,38E-03 |
| Energy dependent regulation of mTOR by LKB1-AMPK | 0,0035 | 28 | 2 | 8,18E-03 |
| Regulation of TP53 Expression and Degradation | 0,0035 | 28 | 2 | 8,18E-03 |
| Lipid digestion, mobilization, and transport | 0,0113 | 91 | 3 | 9,66E-03 |
| TGF-beta receptor signaling activates SMADs | 0,004 | 32 | 2 | 0,0106 |
| Ca2+ pathway | 0,0047 | 38 | 2 | 0,0146 |
| mTOR signalling | 0,0048 | 39 | 2 | 0,0153 |
| PKB-mediated events | 0,005 | 40 | 2 | 0,0161 |
| Heparan sulfate/heparin (HS-GAG) metabolism | 0,0054 | 44 | 2 | 0,0192 |
| NoRC negatively regulates rRNA expression | 0,0061 | 49 | 2 | 0,0235 |
| Class B/2 (Secretin family receptors) | 0,0061 | 49 | 2 | 0,0235 |
| Cyclin B2 mediated events | 0,0006 | 5 | 1 | 0,0239 |
| CREB phosphorylation through the activation of Adenylate Cyclase | 0,0006 | 5 | 1 | 0,0239 |
| Vitamins | 0,0006 | 5 | 1 | 0,0239 |
| Regulation of gene expression in endocrine-committed (NEUROG3+) progenitor cells | 0,0006 | 5 | 1 | 0,0239 |
| Activation of RAS in B cells | 0,0006 | 5 | 1 | 0,0239 |
| Reelin signalling pathway | 0,0006 | 5 | 1 | 0,0239 |
| Negative epigenetic regulation of rRNA expression | 0,0064 | 52 | 2 | 0,0262 |
| IRS-mediated signalling | 0,0309 | 250 | 4 | 0,0319 |
| Insulin receptor signalling cascade | 0,0313 | 253 | 4 | 0,0331 |
| Purine ribonucleoside monophosphate biosynthesis | 0,0009 | 7 | 1 | 0,0332 |
| FGFR3b ligand binding and activation | 0,0009 | 7 | 1 | 0,0332 |
| Opioid Signalling | 0,0077 | 62 | 2 | 0,0362 |
| SHC-related events triggered by IGF1R | 0,0011 | 9 | 1 | 0,0425 |
| Signaling by Insulin receptor | 0,0343 | 277 | 4 | 0,0439 |
| Lipoprotein metabolism | 0,0085 | 69 | 2 | 0,0439 |
| MET activates RAP1 and RAC1 | 0,0012 | 10 | 1 | 0,0472 |

| Reactome Pathway | Ratio of Protein In Pathway | Number of Protein In Pathway | Protein from Gene Set | P-value |
|---|---|---|---|---|
| WNT5A-dependent internalization of FZD2, FZD5 and ROR2 | 0,0012 | 10 | 1 | 0,0472 |
| Adenylate cyclase activating pathway | 0,0012 | 10 | 1 | 0,0472 |
| Cohesin Loading onto Chromatin | 0,0012 | 10 | 1 | 0,0472 |
| E2F-enabled inhibition of pre-replication complex formation | 0,0012 | 10 | 1 | 0,0472 |
| Signaling by TGF-beta Receptor Complex | 0,009 | 73 | 2 | 0,0485 |
| **miR-192-5p** | | | | |
| Hormone-sensitive lipase (HSL)-mediated triacylglycerol hydrolysis | 0.002 | 16 | 4 | 1.78E-03 |
| NGF signalling via TRKA from the plasma membrane | 0.0422 | 341 | 21 | 2.98E-03 |
| Signaling by NODAL | 0.0024 | 19 | 4 | 3.30E-03 |
| Glyoxylate metabolism and glycine degradation | 0.0025 | 20 | 4 | 3.95E-03 |
| Signalling by NGF | 0.0521 | 421 | 24 | 4.06E-03 |
| Regulation of signaling by NODAL | 0.0012 | 10 | 3 | 4.11E-03 |
| Negative regulation of FGFR2 signaling | 0.0042 | 34 | 5 | 4.80E-03 |
| Branched-chain amino acid catabolism | 0.0015 | 12 | 3 | 6.78E-03 |
| Activation of ATR in response to replication stress | 0.0046 | 37 | 5 | 6.79E-03 |
| Signaling by FGFR3 | 0.005 | 40 | 5 | 9.30E-03 |
| Integration of energy metabolism | 0.0114 | 92 | 8 | 9.52E-03 |
| Vasopressin regulates renal water homeostasis via Aquaporins | 0.0032 | 26 | 4 | 9.76E-03 |
| Regulation of pyruvate dehydrogenase (PDH) complex | 0.0017 | 14 | 3 | 0.0103 |
| Signaling by FGFR4 | 0.0051 | 41 | 5 | 0.0103 |
| Ca-dependent events | 0.0035 | 28 | 4 | 0.0125 |
| Negative regulation of FGFR3 signaling | 0.0036 | 29 | 4 | 0.014 |
| CRMPs in Sema3A signaling | 0.002 | 16 | 3 | 0.0147 |
| PKA activation | 0.002 | 16 | 3 | 0.0147 |
| Rap1 signalling | 0.002 | 16 | 3 | 0.0147 |
| Spry regulation of FGF signaling | 0.002 | 16 | 3 | 0.0147 |
| Aquaporin-mediated transport | 0.0037 | 30 | 4 | 0.0157 |
| PKA activation in glucagon signalling | 0.0021 | 17 | 3 | 0.0172 |
| PKA-mediated phosphorylation of CREB | 0.0021 | 17 | 3 | 0.0172 |
| Negative regulation of FGFR4 signaling | 0.0038 | 31 | 4 | 0.0175 |
| Activation of the pre-replicative complex | 0.004 | 32 | 4 | 0.0194 |
| TGF-beta receptor signaling activates SMADs | 0.004 | 32 | 4 | 0.0194 |
| Regulation of insulin secretion | 0.0061 | 49 | 5 | 0.0205 |
| Signaling by FGFR1 | 0.0061 | 49 | 5 | 0.0205 |
| Apoptotic factor-mediated response | 0.0009 | 7 | 2 | 0.021 |
| Glucagon signaling in metabolic regulation | 0.0041 | 33 | 4 | 0.0214 |
| NCAM1 interactions | 0.0041 | 33 | 4 | 0.0214 |
| Negative regulation of FGFR1 signaling | 0.0041 | 33 | 4 | 0.0214 |
| Mitochondrial biogenesis | 0.0064 | 52 | 5 | 0.0256 |
| Signaling by FGFR2 | 0.0088 | 71 | 6 | 0.0263 |
| Lipid digestion. mobilization. and transport | 0.0113 | 91 | 7 | 0.0267 |
| FGFR2b ligand binding and activation | 0.001 | 8 | 2 | 0.0269 |

| Reactome Pathway | Ratio of Protein In Pathway | Number of Protein In Pathway | Protein from Gene Set | P-value |
|---|---|---|---|---|
| Ligand-independent caspase activation via DCC | 0.001 | 8 | 2 | 0.0269 |
| MAPK1 (ERK2) activation | 0.001 | 8 | 2 | 0.0269 |
| Axon guidance | 0.0589 | 476 | 23 | 0.029 |
| Nuclear Events (kinase and transcription factor activation) | 0.0026 | 21 | 3 | 0.0296 |
| Signal transduction by L1 | 0.0026 | 21 | 3 | 0.0296 |
| Signaling by TGF-beta Receptor Complex | 0.009 | 73 | 6 | 0.0296 |
| Signaling by PDGF | 0.0406 | 328 | 17 | 0.0325 |
| Glucagon-like Peptide-1 (GLP1) regulates insulin secretion | 0.0027 | 22 | 3 | 0.0333 |
| SHC-related events triggered by IGF1R | 0.0011 | 9 | 2 | 0.0334 |
| Downstream signal transduction | 0.0379 | 306 | 16 | 0.035 |
| Nuclear signaling by ERBB4 | 0.0028 | 23 | 3 | 0.0372 |
| SHC-mediated cascade:FGFR2 | 0.0028 | 23 | 3 | 0.0372 |
| Signaling by BMP | 0.0028 | 23 | 3 | 0.0372 |
| MyD88 cascade initiated on plasma membrane | 0.0097 | 78 | 6 | 0.0388 |
| Toll Like Receptor 10 (TLR10) Cascade | 0.0097 | 78 | 6 | 0.0388 |
| Toll Like Receptor 5 (TLR5) Cascade | 0.0097 | 78 | 6 | 0.0388 |
| Cohesin Loading onto Chromatin | 0.0012 | 10 | 2 | 0.0404 |
| Interleukin-6 signaling | 0.0012 | 10 | 2 | 0.0404 |
| MAP kinase activation in TLR cascade | 0.0073 | 59 | 5 | 0.0405 |
| PI Metabolism | 0.0073 | 59 | 5 | 0.0405 |
| Signaling by ERBB4 | 0.0052 | 42 | 4 | 0.0453 |
| Signaling by PTK6 | 0.0076 | 61 | 5 | 0.0455 |
| Signaling by EGFR | 0.0392 | 317 | 16 | 0.0455 |
| FRS-mediated FGFR2 signaling | 0.0031 | 25 | 3 | 0.0456 |
| Pyruvate metabolism | 0.0031 | 25 | 3 | 0.0456 |
| L1CAM interactions | 0.0102 | 82 | 6 | 0.0473 |
| Opioid Signalling | 0.0077 | 62 | 5 | 0.0482 |
| Signaling by ERBB2 | 0.0053 | 43 | 4 | 0.0486 |
| Transcriptional activation of mitochondrial biogenesis | 0.0053 | 43 | 4 | 0.0486 |
| MAPK family signaling cascades | 0.0303 | 245 | 13 | 0.0491 |
| Signalling to ERKs | 0.0274 | 221 | 12 | 0.0498 |
| **miR-574-3p** | | | | |
| Dopamine clearance from the synaptic cleft | 0,0002 | 2 | 1 | 1,24E-03 |
| Na+/Cl- dependent neurotransmitter transporters | 0,0007 | 6 | 1 | 3,71E-03 |
| Neurotransmitter Clearance In The Synaptic Cleft | 0,0007 | 6 | 1 | 3,71E-03 |
| WNT5A-dependent internalization of FZD4 | 0,0012 | 10 | 1 | 6,17E-03 |
| Regulation of signaling by NODAL | 0,0012 | 10 | 1 | 6,17E-03 |
| WNT5A-dependent internalization of FZD2, FZD5 and ROR2 | 0,0012 | 10 | 1 | 6,17E-03 |
| Retrograde neurotrophin signalling | 0,0014 | 11 | 1 | 6,79E-03 |
| VLDLR internalisation and degradation | 0,0015 | 12 | 1 | 7,41E-03 |
| Signaling by Activin | 0,0016 | 13 | 1 | 8,02E-03 |
| Signaling by NODAL | 0,0024 | 19 | 1 | 0,0117 |

| Reactome Pathway | Ratio of Protein In Pathway | Number of Protein In Pathway | Protein from Gene Set | P-value |
|---|---|---|---|---|
| LDL-mediated lipid transport | 0,0025 | 20 | 1 | 0,0123 |
| VLDL interactions | 0,0028 | 23 | 1 | 0,0142 |
| Signaling by BMP | 0,0028 | 23 | 1 | 0,0142 |
| Recycling pathway of L1 | 0,0028 | 23 | 1 | 0,0142 |
| Lysosome Vesicle Biogenesis | 0,0031 | 25 | 1 | 0,0154 |
| Golgi Associated Vesicle Biogenesis | 0,0036 | 29 | 1 | 0,0178 |
| Transport of glucose and other sugars, bile salts and organic acids, metal ions and amine compounds | 0,0053 | 43 | 1 | 0,0263 |
| Clathrin derived vesicle budding | 0,0056 | 45 | 1 | 0,0275 |
| trans-Golgi Network Vesicle Budding | 0,0056 | 45 | 1 | 0,0275 |
| Oxygen-dependent proline hydroxylation of Hypoxia-inducible Factor Alpha | 0,0079 | 64 | 1 | 0,039 |
| Lipoprotein metabolism | 0,0085 | 69 | 1 | 0,042 |
| Regulation of Hypoxia-inducible Factor (HIF) by oxygen | 0,009 | 73 | 1 | 0,0444 |
| Cellular response to hypoxia | 0,009 | 73 | 1 | 0,0444 |
| PCP/CE pathway | 0,0097 | 78 | 1 | 0,0474 |
| L1CAM interactions | 0,0102 | 82 | 1 | 0,0497 |

**Supplementary Table S3 –** Primers and reaction components from RT-qPCR reaction performed to analyzed mRNA levels of target genes from miR-100-5p, miR-192-5p or miR-574-3p after mimic treatment in NP69 cells.

| Target Genes | Primers (5' – 3') | Efficiency ($R^2$)¹. |
|---|---|---|
| **miR-100-5p** | | |
| FZD8 | Fwd: CCTCTTCATCGGCACCATGT / Rev: GGTGTAGAGCACGGTGAACA | 0,9983 |
| SMARCA5 | Fwd: GGTCTTGGCATCAATCTTGCG / Rev: CAAGCCTCCCATAGCCTGAA' | 0,9898 |
| **miR-192-5p** | | |
| PRKAR1A [#56 - Xie et al., 2015] | Fwd: GTTTTCGGTCTCCTTTATCGC / Rev: TGCTCTCGGTGTTCCATAAATC | 0.9892¹ |
| RAB2A [#49 - Luo et al., 2015] | Fwd: AGTTCGGTGCTCGAATGATAAC / Rev: AATACGACCTTGTGATGGAACG | 0.9650¹ |
| **miR-574-3p** | | |
| CLTC | Fwd: TGCCATGCCCTATTTCATCCA / Rev: CATCAACTGGGGCTGACCATA | 0,9935 |
| CUL2 | Fwd: ATGTTCTACAGGCTGGTGCG / Rev: TCCTTCCACTGAAATGTTGGCT | 0,9989 |
| EP300 | Fwd: CCGAGACATCTTGAGACGACAG' / Rev: GGGTTGCTGGAACTGGTTATGG | 0,9007 |
| FBXL5 | Fwd: AGCCTCTTTGAAAAGGGACTGA/ Rev: ACATGGGCTGAAAAACCTCCT | 0,9940 |
| PD-L1 | Fwd: GCCCCATACAACAAAATCAACC / Rev: GCTTGTCCAGATGACTTCGG | 0,9971 |
| STC1 | Fwd: AAGATGGCGACCACCAAAGT / Rev: GCAGTGACGCTCATAAGGGA | 0,9926 |
| **Reference genes** | | |
| HSPCB | Fwd: TCTGGGTATCGGAAAGCAAGCC / Rev: GTGCACTTCCTCAGGCATCTTG | 0.9799¹ |
| RPS13 | Fwd: CGAAAGCATCTTGAGAGGAACA / Rev: TCGAGCCCAAACGGTGAATC | 0.9986¹ |
| RRN18S | Fwd: AGAAACGGCTACCACATCCA / Rev: CACCAGACTTGCCCTCCA | 0.9946¹ |

1 Reaction components (GoTaq® qPCR Master Mix 2X. 0.2 µM primers. Nuclease-free water (NF)²) and cycling protocol (95°C - 2 min (1x); 95°C - 15s. 60°C - 60s. 72°C – 30s (40x); 60-95°C (1x)³) was performed according to the manufacturer instructions.

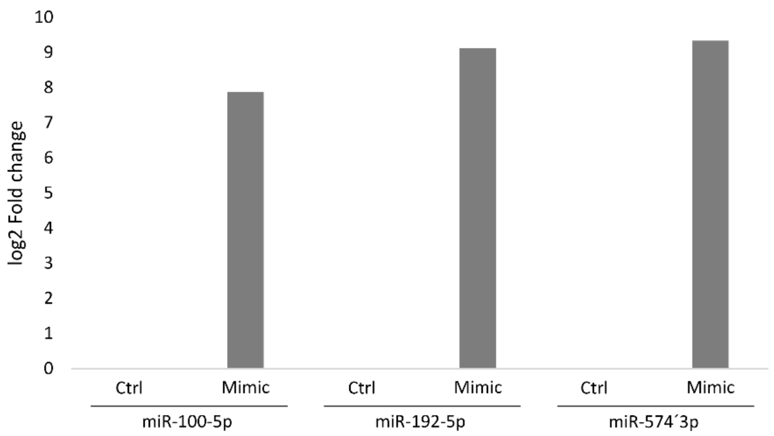


Supplementary Figure S2 – Relative expression of miR-100-5p, miR-192-5p, and miR-574-3p in NP69 cells 24 h after transfection with 10nM of the respective mimics compared to control (reagent only).
